# A multilayer network model of neuron-astrocyte populations in vitro reveals mGluR_5_ inhibition is protective following traumatic injury

**DOI:** 10.1101/798611

**Authors:** Margaret E. Schroeder, Danielle S. Bassett, David F. Meaney

**Affiliations:** Department of Bioengineering, School of Engineering & Applied Science, University of Pennsylvania, Philadelphia, PA, USA; Department of Physics & Astronomy, College of Arts & Sciences, University of Pennsylvania, Philadelphia, PA, USA; Department of Electrical & Systems Engineering, School of Engineering & Applied Science, University of Pennsylvania, Philadelphia, PA, USA; Department of Neurology, Perelman School of Medicine, University of Pennsylvania, Philadelphia, PA, USA; Department of Psychiatry, Perelman School of Medicine, University of Pennsylvania, Philadelphia, PA, USA

**Author notes:** Corresponding author: David F. Meaney.

**Keywords:** Astrocyte, Traumatic Injury, Multilayer Network, Neuron-glia Signaling

## Abstract

Astrocytes communicate bidirectionally with neurons, enhancing synaptic plasticity and promoting the synchronization of neuronal microcircuits. Despite recent advances in understanding neuron-astrocyte signaling, little is known about astrocytic modulation of neuronal activity at the population level, particularly in disease or following injury. We used high-speed calcium imaging of mixed cortical cultures *in vitro* to determine how population activity changes after disruption of glutamatergic signaling and mechanical injury. We constructed a multilayer network model of neuron-astrocyte connectivity, which captured distinct topology and response behavior from single cell type networks. mGluR_5_ inhibition decreased neuronal activity, but did not on its own disrupt functional connectivity or network topology. In contrast, injury increased the strength, clustering, and efficiency of neuronal but not astrocytic networks, an effect that was not observed in networks pre-treated with mGluR_5_ inhibition. Comparison of spatial and functional connectivity revealed that functional connectivity is largely independent of spatial proximity at the microscale, but mechanical injury increased the spatial-functional correlation. Finally, we found that astrocyte segments of the same cell often belong to separate functional communities based on neuronal connectivity, suggesting that astrocyte segments function as independent entities. Our findings demonstrate the utility of multilayer network models for characterizing the multiscale connectivity of two distinct but functionally dependent cell populations.

**AUTHOR SUMMARY:** Astrocytes communicate bidirectionally with neurons, enhancing synaptic plasticity and promoting the synchronization of neuronal microcircuits. We constructed a multilayer network model of neuron-astrocyte connectivity based on calcium activity in mixed cortical cultures, and used this model to evaluate the effect of glutamatergic inhibition and mechanical injury on network topology. We found that injury increased the strength, clustering, and efficiency of neuronal but not astrocytic networks, an effect that was not observed in injured networks pre-treated with a glutamate receptor antagonist. Our findings demonstrate the utility of multilayer network models for characterizing the multiscale connectivity of two distinct but functionally dependent cell populations.

## INTRODUCTION

Astrocytes, the most abundant glial cell in the brain, can modulate the activity of neurons through the uptake of neurotransmitters and release of gliotransmitters (Di Castro et al. (2011); J. Kang, Jiang, Goldman, and Nedergaard (1998); Panatier et al. (2011); Perea and Araque (2007); J. T. Porter and McCarthy (1996)). One such pathway is uptake of synaptic glutamate through the astrocytic metabotropic glutamatergic receptor subtype 5 (mGluR_5_) (Panatier and Robitaille (2016), Fig. 1. Through gliotransmission, astrocytes can enhance or depress synaptic plasticity and the synchronization of neuronal circuits (Pascual et al. (2005); Poskanzer and Yuste (2011); Serrano, Haddjeri, Lacaille, and Robitaille (2006)). Although prior studies disagree on the relative abundance of mGluR_5_ in neurons vs. astrocytes Kettenmann and Schachner (1985); Saunders et al. (2018); Zhang et al. (2014), astrocyte mGluR_5_ has been identified as one of the most prominent receptors implicated in astrocytic detection of neurotransmitters, Ca2+ signaling, and downstream synaptic modulation (Panatier and Robitaille (2016); Panatier et al. (2011); J. T. Porter and McCarthy (1996); X. Wang et al. (2006)). Despite recent advances in understanding neuron-astrocyte signaling, comparatively little is known about astrocytic modulation of neuronal population activity, particularly in cases where that activity has been altered by disease and by injury. In this study, we sought to define the role of astrocytes in modulating neural circuits before and after targeted neuronal injury.

**Figure 1.**
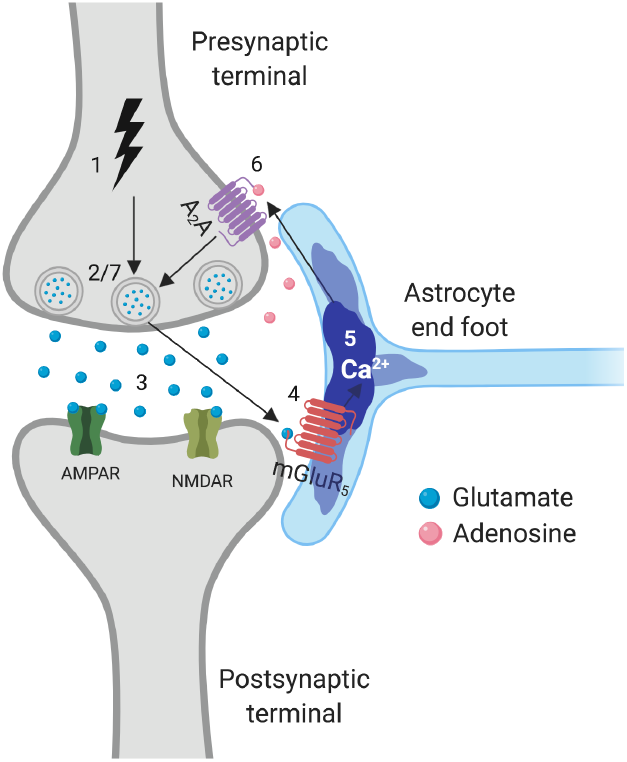
The Tripartite Synapse mediated through mGluR_5_. An action potential arrives at the presynaptic terminal (1) and triggers glutamate release (2-3). Glutamate acts on astrocytic mGluR_5_ (4), leading to an increase in astrocyte Ca^2+^ (5). The astrocyte releases adenosine (6), which is taken up by presynaptic A_2_A receptors, and alters the probability of neurotransmitter release (7). Created with BioRender.

Based on the large body of evidence supporting a role for astrocytes in establishing neural population synchronization and in protecting against neural degeneration following traumatic brain injury (TBI) (Burda, Bernstein, and Sofroniew (2016); Myer, Gurkoff, Lee, Hovda, and Sofroniew (2006)), astrocytes may play a key role in recovering neuronal network synchrony, which is disrupted following moderate mechanical injury to neural tissue (W. H. Kang et al. (2015); Patel, Ventre, and Meaney (2012)). Previous studies examining astrocyte response to mechanical stretch injury *in vitro* directly triggered injury cascades in astrocytes (Charles, Merrill, Dirksen, and Sandersont (1991); Ellis, McKinney, Willoughby, Liang, and Povlishock (1995); Rzigalinski, Weber, Willoughby, and Ellis (1998)) and reactive gliosis (Miller et al. (2009)). Mediated by intracellular calcium waves in astrocytes that originate from sites of mechanical injury, past work shows signaling from astrocytes can temporarily silence neuronal network after trauma (Choo et al. (2013)). In addition, therapeutically manipulating the receptors that control the extent of astrocyte intercellular waves after trauma also lead to an improvement in outcome after injury. However, the interdependence of targeted neuronal injury and astrocyte activity, and how these cell types interact to recover activity in neuronal activity in networks, is not known. Determining more precisely the nature of neuron-astrocyte communication following injury may reveal mechanisms on how neural circuits recover from trauma, and the role that surrounding astrocyte networks play in this recovery process.

*In vivo*, the stress transferred to the cellular and subcellular components is complex and heterogeneous (Bain, Shreiber, and Meaney (2003); Bar-Kochba, Scimone, Estrada, and Franck (2016); Cloots, Van Dommelen, Nyberg, Kleiven, and Geers (2011); LaPlaca, Cullen, McLoughlin, and Cargill II (2005); Pan, Sullivan, Shreiber, and Pelegri (2013)). Rather than applying a uniform membrane deformation to a mixture of integrated neuronal and glial cells in a network, we sought to mimic the heterogeneous nature of mechanical trauma using a localized injury model developed recently (Mott, von Reyn, Firestein, and Meaney (2021)). One of the primary events that occurs instantly during trauma is the activation of receptors and channels in the membrane (Iwata et al. (2004); Patel, Ventre, Geddes-Klein, Singh, and Meaney (2014)), the instantaneous release of neurotransmitters and gliotransmitters (Choo et al. (2013); LaPlaca and Thibault (1998)), and the appearance of nonspecific pores in the plasma membrane (Farkas, Lifshitz, and Povlishock (2006); Kilinc, Gallo, and Barbee (2009); LaPlaca et al. (2019); Tehse and Taghibiglou (2019)) that can impair neuronal and glial function. Within this injury model, we chose to focus on the relative role of mGluR_5_ receptors after traumatic neuronal injury *in vitro*. The mGluR_5_ receptor is an appealing receptor target because it is expressed by both neurons and astrocytes, yet very little is known about its effect on network activity and structure. Moreover, evidence points to an uncertain role for mGluR_5_ on neuronal outcome. Some studies show that activation of mGluR_5_ improves outcome following injury (Chen, Cao, et al. (2012); Loane, Stoica, Byrnes, Jeong, and Faden (2013); Loane et al. (2014)), while other reports show inhibition is necessary to reduce neuronal degeneration and improve functional recovery (Chen, Zhang, et al. (2012); Movsesyan et al. (2001)). Part of this controversy could be that these studies could not study how the activation of these receptors affects the structure and topology of neuronal microcircuits after trauma. With the aforementioned injury mechanisms occurring at the moment of injury, our approach to determine the mechanistic role of mGluR_5_ on neuronal and astrocyte network structure required us to block mGluR_5_ prior to mechanical trauma.

Past work to study the dynamics between astrocytes and neurons often relies on *in situ* techniques (e.g., acute slice preparations) or transgenic approaches, neither of which allows an exact determination of how neighboring astrocytes influence the activity of neurons, or vice versa. Although the emergence of probes to record cellular-level resolution of neuronal networks *in vitro* and *in vivo* emerged in the past decade (Akerboom et al. (2012); Tian et al. (2009)), the relationship between the timing and influence of astrocytic activation – which often occurs over several seconds – on the subsequent neuronal activity patterns using these probes or other technologies remains largely unknown. One key obstacle is that this relationship depends on both proximity of one cell type to another, the resulting connections each cell types forms within a larger network, and the complex timing of these interactions.

The emerging field of network neuroscience, which uses tools from graph theory to analyze neural communication as reflected in the topology of interaction patterns, provides an apt set of tools for analyzing multicellular activity and, in particular, how this activity can be causally linked across cell populations, or layers. Multilayer networks are a generalized framework for incorporating multiple types of relations between participating units, capable of describing and modeling a wide range of complex systems (Betzel and Bassett (2017); Kivelä et al. (2014); Muldoon and Bassett (2016); Vaiana and Muldoon (2018)). The network formed from bidirectional neuron-astrocyte functional connectivity lends itself to a multilayer network model, where astrocytes and neurons are encoded as nodes in a multilayer network. In analyzing dynamic activity occurring across both neurons and astrocytes within the framework of a single network, one can objectively assess the relative influence of one cell type on another, understand whether these cell layers communicate with each other, and identify the role of exogenous factors, such as mGluR_5_ signaling and traumatic injury, on network structure and function.

The objective of this study was threefold: to determine the spatial and functional topology of neuron-astrocyte networks using a multilayer network model, to characterize changes in topology resulting from single-cell injury and mGluR_5_ inhibition, and to determine whether fully functional cellular communication is required for post-injury recovery of baseline network topology. We show that targeted neuronal injury, but not pharmacological blockade of glutamatergic signaling via mGluR_5_ altered higher-order topological properties of multilayer networks, including clustering and global efficiency of the neuronal layer. Our results also indicate that mGluR_5_ inhibition modulates the topological impact of injury. More broadly, our findings demonstrate the utility of multilayer network models for characterizing the micro-, meso-, and global-scale connectivity of two morphologically and genetically distinct but functionally dependent cell populations.

## RESULTS

We used a multilayer and two single-layer network models to characterize population-level neuron and astrocyte segment dynamics. Neural cell cultures can be modeled as networks without violating the foundational modeling assumptions of network science (Bassett, Zurn, and Gold (2018)). A node represents a single neuron in the neural network, and it represents an astrocyte process in the astrocyte network. We use moderate speed (*>* 10 frames per second) imaging of intracellular calcium in individual cells to detect transient events in these network nodes. In neurons, these transient changes in fluorescence represent one more action potential event (Akerboom et al. (2012); Tian et al. (2009)), which can release neurotransmitters that can activate surface receptors on astrocytes to subsequently release of calcium from intracellular stores (Araque, Parpura, Sanzgiri, and Haydon (1999)). Likewise, calcium transients in astrocytes can release gliotransmitters which influence neuronal activity (Araque, Parpura, Sanzgiri, and Haydon (1998); Parpura and Haydon (2000)). Therefore, tracking the timing of fluorescent changes in each cell types provides a rich data set containing the spatial location and temporal activity of each cell in the population. Edges are determined by functional connectivity between nodes based on fluorescence activity over time (Patel, Man, Firestein, and Meaney (2015)). Each node was assigned to the neuron or astrocyte layer depending on its identity (Fig. 2, SI Sec. B). Each edge representing a functional connection between an astrocyte and a neuron, or between two astrocytes or two neurons, was weighted by the functional connectivity between them (see Methods, SI Sec. C).

**Figure 2.**
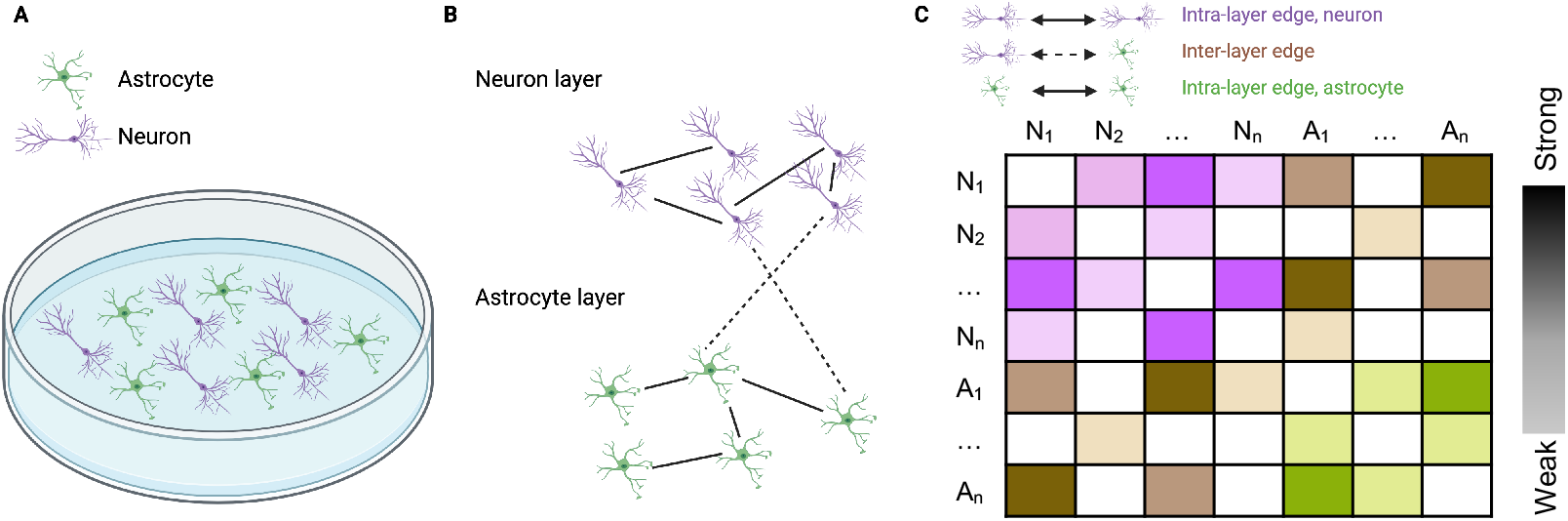
A multilayer network model of neurons and astrocytes in co-culture. **A**. Simplified schematic of neurons (purple) and astrocytes (green) in co-culture. Morphology and size are exaggerated. **B**. Cells are separated into two distinct layers based on their identity. Layers are made up of exclusively one cell type. Each cell may be connected to cells in its own layer or to cells in the other layer. Intralayer edges are depicted with full lines and interlayer edges with dashed lines. **C**. Example adjacency matrix for the multilayer network shown in panel **B**. The purple and green weights correspond to intralayer edges and the brown weights correspond to interlayer edges. Color saturation indications strength of connection.

Only cells that were inactive across all conditions were eliminated from the graph, allowing some nodes to have no edges in certain conditions. Dishes were analyzed as separate networks, each with a variable number of nodes (Fig. S1C). Many useful statistics can be calculated from these graphs, the simplest being the number of nodes and edges. More biologically informative, however, are network parameters describing how cells are functionally connected. Because the plating density of neurons and astrocytes varies between dishes and between isolations, we calculated network measures that can be normalized for network size (SI Sec. D). Mean degree, density, nodal strength (normalized to network size), clustering coefficient, betweenness centrality (also normalized), and global efficiency were calculated for each network and for a random null network model in which both degree distribution and density were preserved. The rationale for reporting these metrics was biologically driven, as they capture general properties of multicellular communication: its overall density, the extent to which connected cells signal to shared neighbors, and the extent to which cells act as hubs, facilitating signaling between other cells. A random null model was chosen because the attachment of cortical neurons and astrocytes to the dish upon plating was random, as was their wiring to other cells after plating.

### Blockade of glutamatergic signaling and mechanical injury decrease the activity of neurons, but not astrocytes

As a first-order measure of network activity, we calculated the mean frequency of calcium events for each cell type in each dish (SI Sec. A, SI Sec. B), and then averaged these values over all cells in the field of view that were active at any time point during the experiment. Neuronal event rate decreased following treatment with 2-Methyl-6-(phenylethynyl)pyridine (MPEP), a potent mGluR_5_ inhibitor, immediately after drug addition but before injury. Following injury, neuronal event rate was lower in the MPEP-treated and injured arms relative to untreated control dishes (Fig. S1A). However, there was no significant difference in event rate between baseline and post-injury time points for any of the four groups. In the untreated, uninjured control group (MEM, Sham) and the MPEP-treated uninjured group, there was a decrease in event rate following drug treatment, followed by a subsequent increase to near-baseline levels or higher at the injury time point. This suggests that the addition of vehicle depresses event rate, which is subsequently recovered in uninjured networks. Both mGluR_5_ blockade and neuronal injury significantly reduced neuronal event rate relative to untreated dishes (Fig. S1A and Table S1; Tukey’s multiple comparisons test following 2-way ANOVA, at post-injury time-point, *p* = 0.0111 for MEM, Sham vs. MEM, Injury, *p* = 0.0145 for MEM, Sham vs. MPEP, Sham, *p* = 0.0004 for MEM, Sham vs. MPEP, Injury). Astrocytic event rate was unchanged following neuronal injury and/or mGluR_5_ inhibition (Fig. S1B, Table S1). These results suggest that any changes in functional network topology may be mediated directly through decreased activity in neural layers, while changes in astrocyte layer topology are unlikely to be mediated through decreased astrocyte activity.

### Non-random topology of neuron, astrocyte, and multilayer networks

Next, we sought to characterize functional network topology, which is a higher-order measure of coherent population-level activity. Weighted adjacency matrices were generated by calculating the Pearson’s correlation coefficient between pairs of filtered and scaled astrocyte calcium traces (SI Sec. C). In addition to the network density *κ*, the global efficiency *E*, the clustering coefficient *C*, the normalized degree *K*, and the normalized betwenness centrality *B*, we calculated the mean normalized nodal strength *S* of each network and normalized it based on the number of nodes in the network. To test whether these network statistics were topologically meaningful (non-random) after controlling for network size and density, we compared them to those of a null model in which connections were randomly rewired while preserving the degree distribution and density. Relative to randomized networks, neuron networks exhibited significantly greater clustering coefficient (Fig. 3A; one-sample *t*-test on differences between actual values and null model values of *C, t* = 6.010, *df* = 35, *p <* 0.0001), significantly lower betweenness centrality (one-sample *t*-test on differences between actual values and null model values of *B, t* = 2.839, *df* = 35, *p* = 0.0075) and significantly lower global efficiency (one-sample *t*-test on differences between actual values and null model values of *E, t* = 9.518, *df* = 35, *p <* 0.0001). That is, neuron networks have more interconnected triads and fewer long-distance connections, which promote efficiency of communication between nodes, than would be expected based purely on their density. As would be expected for random networks of the same size and degree distribution, the clustering coefficient *C* and global efficiency *E* were strongly correlated with edge density, while betweenness centrality *B* was anti-correlated (Fig. 3B; see Table SX for statistics). The degree to which these topological measures depended on density in our observed networks was slightly different than that of randomized control networks (Fig. S2, Table S2).

**Figure 3.**
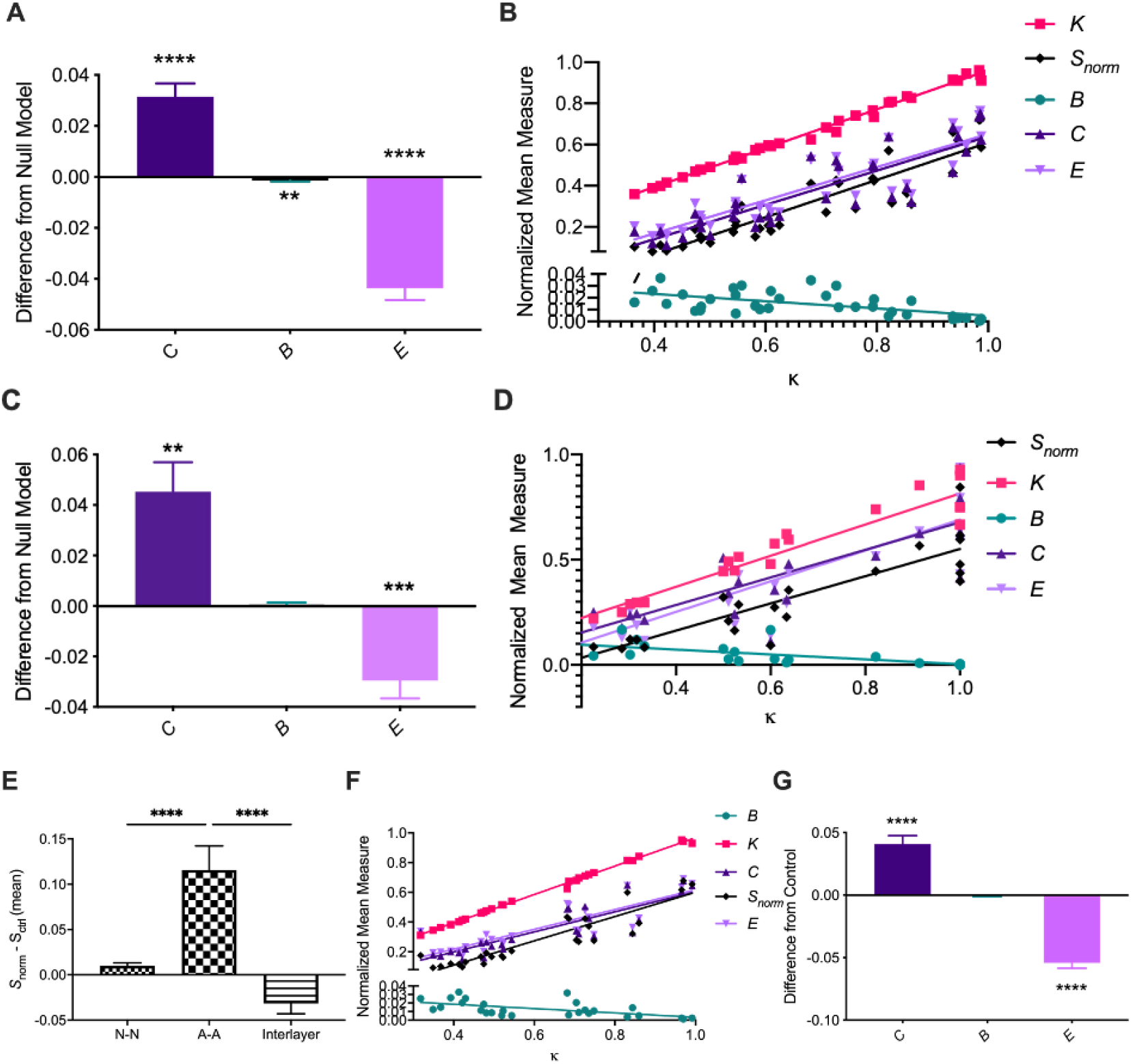
Characterization of neuron-neuron, astrocyte-astrocyte, and multilayer neuron-astrocyte functional network topology **A**. Difference from random null model of calculated mean clustering coefficient *C*, normalized betweenness centrality *B*, and global efficiency *E* for neuron-neuron networks. We observe significantly larger clustering coefficients and significantly lower global efficiency than expected from a random null model. **B**. Mean clustering coefficient *C*, normalized betweenness centrality *B*, normalized degree *K*, normalized strength *S*_*norm*_, and global efficiency *E* vs. mean density *κ* for each dish at the third imaging time point (1 hour post-injury). We observe clear positive correlations as assessed by a linear regression for *K, S*_*norm*_, *C*, and *E*, and a clear negative correlation for *B*. **C**. Same as **A**, for astrocyte-astrocyte networks. A similar network topology is observed. **D**. Same as **B**, for astrocyte-astrocyte networks. **E** Difference in mean normalized strength of neuron layer, astrocyte layer, interlayer, and multilayer connections, compared to a randomized network with preserved degree distribution and density. **F**. Same as **B**, for multilayer networks. **G**. Same as **A**, for multilayer networks. Error bars indicate standard error of the mean (SEM) and asterisks indicate statistical significance (*p *≤* 0.05, **p *≤* 0.01, ***p *≤* 0.001, ****p *≤* 0.0001).

In general, astrocyte networks were small and sparse. After eliminating segments that were not active at any time during the experiment, only 28 out of 36 dishes remained, with a median of about 3 active whole cells and about 15 active astrocyte processes per dish (Fig. S1C, Fig. S3D). The number of whole astrocytes correlated well with the number of active astrocyte segments, with each active cell having on average around five active segments (Fig. Fig. S3D). Similar to the neuron networks, functional astrocyte networks displayed significantly higher clustering coefficient (one-sample *t*-test on differences between the actual values and null model values of *C, t* = 3.887, *df* = 21, *p* = 0.0009) and significantly lower global efficiency (one-sample *t*-test on differences between actual values and null model values of *E, t* = 4.167, *df* = 21, *p* = 0.0004) than control networks (Fig. 3C). For astrocyte networks, *B* was not significantly different from that of randomized control networks (one-sample *t*-test on differences between actual values and null model values of *B, t* = 0.6922, *df* = 21, *p* = 0.4964). As for neuron networks, *C* and *E* were strongly correlated with edge density for astrocyte networks, while betweenness centrality *B* was anti-correlated (Fig. 3D; see Table S2 for statistics). Again, the degree to which these topological measures depended on density in our observed networks was slightly different than that of randomized control networks (Fig. S2, Table S2).

While it is useful to consider neuron and astrocyte networks separately, neurons and astrocytes do not exist in isolation. To better understand the interactions between neurons and astrocytes, and to assess whether the topology of these sub-populations is independent of or informed by the other, we modeled neuron-astrocyte functional interactions as a multilayer network (Fig. 2). Edges for multilayer adjacency matrices were assigned weights from the functional connectivity between neurons and neurons, between astrocytes and astrocytes, and between neurons and astrocytes, as measured by the Pearson’s correlation coefficient of filtered and scaled calcium traces (SI Sec. C). Strength and density were calculated for the entire multilayer network, for neuron and astrocyte layers separately, and for interlayer connections separately. Mean normalized betweenness centrality, mean clustering coefficient, and global efficiency were calculated for the entire multilayer network and for control networks generated as previously described.

Nodal strength in the neuron network layer (Fig. 3E, one-sample *t*-test on differences between actual values and null model values of *S, t* = 2.913, *df* = 26, *p* = 0.0073) and in the astrocyte network layer (one-sample *t*-test on differences between actual values and null model values of *S, t* = 4.338, *df* = 22, *p* = 0.0003) was higher than expected in randomized multilayer networks that preserved strength. Interestingly, mean nodal strength calculated only from interlayer edges was lower than that expected in randomized controls (one-sample *t*-test on differences between actual values and null model values of *S, t* = 2.716, *df* = 26, *p* = 0.0104). Further, the astrocyte layer nodal strength was significantly larger than that of the neuron layer (Tukey’s multiple comparisons test following ordinary one-way ANOVA, *q* = 6.692, *df* = 74, *p <* ¡0.0001) and than that of interlayer edges (Tukey’s multiple comparisons test following ordinary one-way ANOVA, *q* = 9.313, *df* = 74, *p <* ¡0.0001). In other words, intralayer connections are stronger than interlayer connections, and astrocyte segments are more strongly interconnected than neurons, compared to random networks maintaining the same mean nodal strength.

Consistent with our findings for neuron and astrocyte networks individually, multilayer networks exhibit significantly larger clustering coefficient *C* (one-sample *t*-test on differences between actual values and null model values, *t* = 6.190, *df* = 26, *p <* 0.0001) and lower global efficiency *E* (one-sample *t*-test on differences between actual and null, *t* = 12.27, *df* = 26, *p <* 0.0001) than their randomized counterparts (Fig. 3G). No differences in betweenness centrality *B* were observed (one-sample *t*-test on differences between actual and null, *t* = 1.095, *df* = 26, *p* = 0.2835). Also consistent with our findings for the individual layers, *C* and *E* were strongly correlated with edge density for multilayer networks, while betweenness centrality *B* was anti-correlated (Fig. 3F, Table S2). Again, the degree to which these topological measures depended on density in our observed networks was slightly different than that of randomized control networks (Fig. S2, Table S2).

### Mechanical injury alters neuron and multilayer, but not astrocyte layer network topology

Because glutamate is an important neural and glial transmitter, we targeted mGluR_5_ to inhibit neuron-to-neuron (N-N) and neuron-to-astrocyte (N-A) communication. We then measured the resulting changes in network topology by calculating the mean nodal strength (*S*), density (*κ*), clustering coefficiency (*C*), betweenness centrality (*B*), and global efficiency (*E*) of functional neuronal networks after disruption of mGluR_5_-mediated N-N and N-A signaling (SI Sec. D). Importantly, for both neurons and astrocytes, mean nodal strength and edge density were not positively correlated with mean event rate (Fig. S4A, Fig. S5A) Thus, any observed changes in network density and/or topology from our manipulations are likely not caused by spurious changes in calcium trace correlation due to reduced or increased activity.

Both administration of the mGluR_5_ antagonist and neuronal injury led to a modest increase in mean edge density and mean nodal strength at the final experimental time point, though this different was not statistically significant (Fig. S4B-D, one-way ANOVA between treatment groups for *S*_*norm*_ 1 hour post-injury, *F* (3, 32) = 1.741, *p* = 0.1784). To assess the impact of mGluR_5_ inhibition, injury, and their interaction on multilayer network topology, we fit a generalized linear model to multilayer graph statistics including the betweenness centrality *B*, the clustering coefficient *C*, and the global efficiency *E*. The outcome variable of interest was calculated for each dish at the third time point and regressed on its group assignment while controlling for mean nodal strength as a covariate of non-interest. We found that mean nodal strength was the greatest, and most statistically significant, covariate of all measures (Table S3). The dependence of these topological properties on density is also evidenced by a strong correlation between the density and the clustering coefficient *C*, the global efficiency *E*, and the betweenness centrality *B* (Fig. 3B). At the final experimental time point, mean nodal strength *S* of multilayer networks was significantly altered by injury only (GLM with *S* as outcome variable and MPEP, Injury, MPEP + Injury interaction term, and event rate as covariates; two-sided *z*-test on estimated coefficient for *S* on Sham dummy variable, *β* = -0.2046, *z* = -2.183, *p* = 0.029). After controlling for mean nodal strength, the GLM revealed that injury increased clustering coefficient *C* and global efficiency *E* of neuronal networks (two-sided *z*-test on estimated coefficient for *C* on Sham dummy variable, *β* = -0.0346, *z* = -2.450, *df* = 31, *p* = 0.014; coefficient on Sham dummy variable for *E, β* = -0.0368, *z* = -2.207, *df* = 31, *p* = 0.027). This effect was not seen in the MPEP + Inj group, suggesting that the addition of MPEP attenuates the effect of injury.

We observed no significant difference in density or mean strength of astrocyte networks after treatment with anti-mGluR_5_ or after injury Fig. S5B-D). As for neuron networks, we used a GLM and regressed out mean nodal strength in astrocyte networks to isolate the effect of our manipulations. For all measures, there was no significant effect of MPEP administration, injury, or their interaction (Table S4). Furthermore, there was no significant effect of MPEP, injury, or their interaction on mean nodal strength of astrocyte networks at the final time point (GLM with *S* as outcome variable and MPEP, Injury, MPEP + Injury interaction term, and event rate as covariates, all *p*-values from *z*-tests on coefficients above 0.05).

Finally, as for the neuron and astrocyte layers, we fit a GLM to isolate the effect of our manipulations on full multilayer topology at the final experimental time point (1 hour post-injury). Unlike the behavior of the neuronal layer, there was no significant effect of MPEP, injury, or their interaction on either mean nodal strength of multilayer networks or inter-layer connections at the final time point (GLM with *S* as outcome variable and MPEP, Injury, MPEP + Injury interaction term, and event rate as covariates).

However, in the injury-only group, but not the MPEP + Injury group, we observed a significant increase in the clustering coefficient *C* (two-sided *z*-test on estimated coefficient on Sham dummy variable for *C, β* = -0.0368, *z* = -2.473, *df* = 22, *p* = 0.013), but no change in the global efficiency *E* or betweenness centrality *B*. Neither *C, B*, or *E* were significantly different from control in MPEP-treated or MPEP-treated and injured groups, suggesting that the effect of injury on multilayer network topology is attenuated by mGluR_5_ inhibition. It is notable that injury had a similar effect in direction and magnitude on *C* for neuronal and multilayer networks, but that the observed increase in *E* for injured neuronal networks was not observed in multilayer networks.

To measure the impact of the relative abundance of neurons to astrocytes on multilayer topology, we sub-sampled the neuron population to equal the number of astrocytes and recalculated network statistics on these synthetic half-neuron, half-astrocyte networks (SI Sec. E). Sub-sampled multilayer networks exhibited similar baseline topology (Fig. S6), but slightly different response behavior (Table S7) than full multilayer networks, suggesting that the relative abundance of neurons partially drives changes in topology.

### Mechanical injury increases the dependence of functional connectivity on spatial proximity in mesoscale multilayer networks

In addition to functional connectivity, spatial distance between nodes might shape network topology (Fig. 4A-B, Fig. S7A-D), though the baseline magnitude of this dependence is likely to be small for out networks, given the spatially random plating of cells in dissociated culture and the scale of the imaging field of view (usually less than 100 cells per field of view). The location of each cell in 2D space was extracted from calcium fluorescence recordings. Spatial adjacency matrices were generated such that nodes with the shortest radial distance were most strongly connected. To test this hypothesis, we regressed the the edge weights in the functional adjacency matrix versus the edge weight for the same node pair in the spatial adjacency matrix for all connection types in our multilayer networks (neuron-neuron, neuron-astrocyte, and astrocyte-astrocyte) at baseline. We found a very small but significant correlation between spatial and functional weights for neuron-neuron and astrocyte-astrocyte layers (simple linear regression; neuron-neuron, 95% CI of slope [0.1463, 0.1823], *R*^2^ = 0.006624, *df* = 48,140, *p <* 0.0001; astrocyte-astrocyte, 95% CI of slope [0.2373, 0.3816], *R*^2^ 0.01501, *df* = 4,643, *p <* 0.0001; neuron-astrocyte, 95% CI of slope [-0.2602, 0.01779], *df* = 20,733, *p* = 0.7127). When measured as a 2-dimensional correlation coefficent (Fig. S7H), the spatial-functional agreement was significantly stronger than for randomized control networks for neuron-neuron and astrocyte-astrocyte connections (95% CI of mean difference for N-N connections, [0.07024, 0.1060]; 95% CI of mean difference for A-A connections, [0.2166, 0.4144]) but weaker for inter-layer neuron-astrocyte connections (95% CI of mean difference for N-A connections, [-0.3153, -0.1516]). The finding that astrocyte-astrocyte functional connectivity was more highly correlated to spatial connectivity than for neuron networks (Tukey’s multiple comparisons test following mixed effects analysis, *q* = 9.415, *df* = 20, *p <* 0.0001) is expected given that microdomains of the same astrocyte were counted as separate nodes.

**Figure 4.**
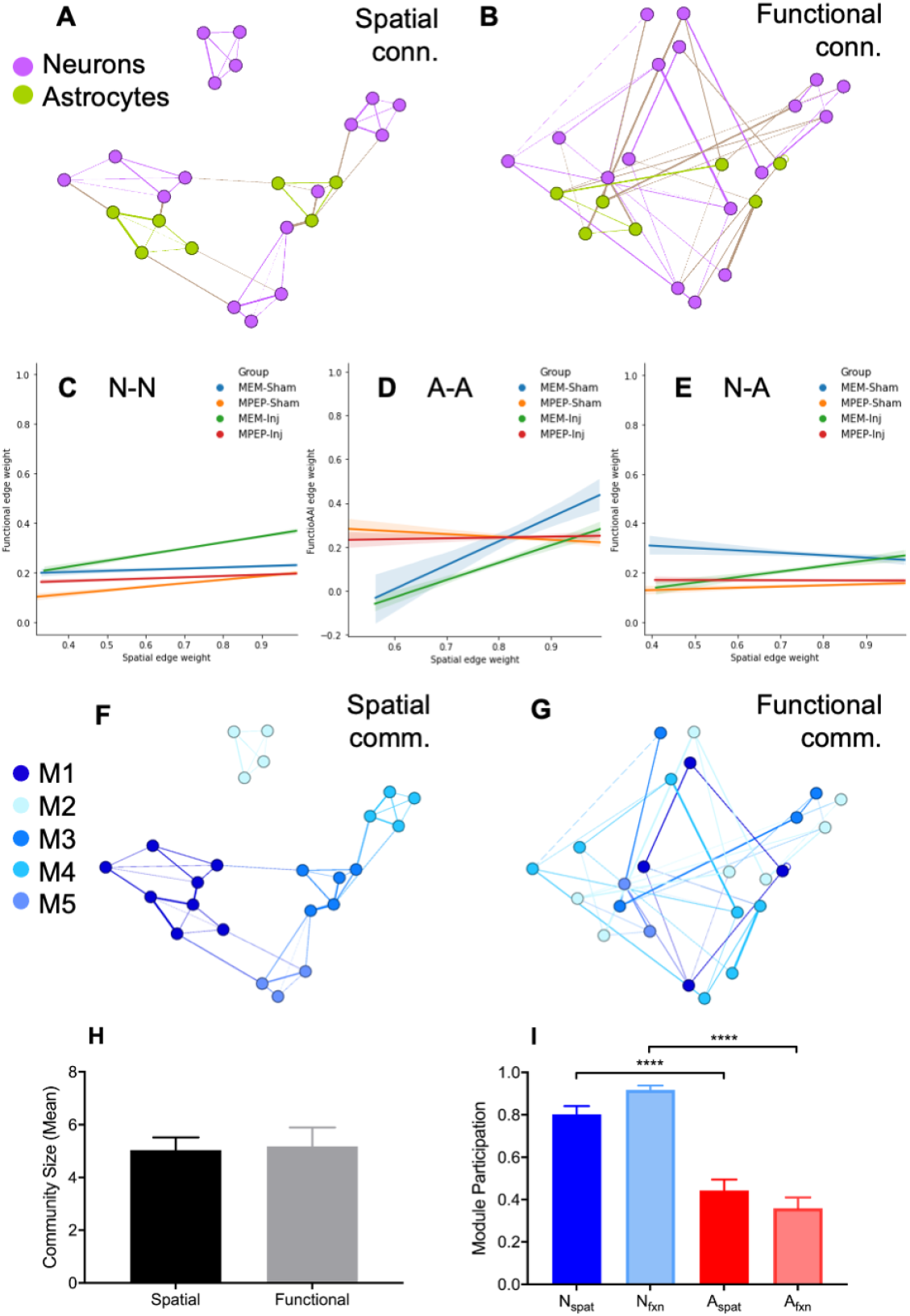
Spatial-functional dependency in multilayer networks. **A**. A representative graph of a multilayer network with weighted edges based on spatial proximity. **B**. A representative graph of a multilayer network with weighted edges based on functional connectivity. **C-E**. Functional edge weight versus spatial edge weight for neuron-neuron (**G**), astrocyte-astrocyte (**H**), and neuron-astrocyte (**H**) layers of the multilayer network. Shown is the best-fit line from simple linear regression for the four experimental groups at the final time point (1 hour post-injury). Shading indicates 95% confidence interval on the slope (Table S8). **F**. Community structure of the network shown in panel **A** with modules determined based on spatial proximity. Nodes of the same color belong to the same spatial module. **G**. Community structure of the network shown in panel **F** with modules determined based on functional connectivity. Nodes of the same color belong to the same functional module. If functional connectivity were based on spatial proximity, the modules in panels **F** and **G** would be the same or highly similar. **H**. Mean community size does not differ significantly between functional and spatial multilayer communities, as the spatial tuning parameter was adjusted to minimize this difference. **I**. Average module participation, the fraction of modules that contain at least one of that cell type, as determined based on spatial distance and functional connectivity for both neurons and astrocytes. Error bars indicate standard error of the mean (SEM) and asterisks indicate statistical significance (*p *≤* 0.05, **p *≤* 0.01, ***p *≤* 0.001, ****p *≤* 0.0001).

If functional neuronal connections are not spatially restricted, as the observed low correlation between spatial and functional connectivity would suggest, neurons and astrocyte microdomains must be able to form relatively long-distance functional connections. Assuming there is both a spatial (albeit small) and non-spatial component influencing functional connectivity, and that the functional, but not spatial, component is disrupted following injury, then mesoscale topology should change to favor connectivity governed by short-distance spatial connections following injury. To test this hypothesis, we compared the correlation between spatial and functional connectivity (weights in the functional or spatial adjacency matrix) at the final experimental time point between different experimental groups for the neuron-neuron, astrocyte-astrocyte, and interlayer connections (Fig. 4). As predicted, we found that injury increased the correlation between spatial and functional connectivity for the neuronal layer (Fig. 4C, simple linear regression, *R*^2^ = 0.0 for MEM, Sham, *R*^2^ = 0.01 for MEM, Inj), astrocyte later (Fig. 4D, simple linear regression, *R*^2^ = 0.075 for MEM, Sham, *R*^2^ = 0.096 for MEM, Inj), and interlayer connections (Fig. 4E, simple linear regression, *R*^2^ = 0.001 for MEM, Sham, *R*^2^ = 0.0007 for MEM, Inj; Table S8). Treatment with MPEP decreased the spatial-functional correlation for the astrocyte layer (simple linear regression, *R*^2^ = 0.075 for MEM, Sham, *R*^2^ = 0.002 for MPEP, Sham), but increased it for the neuronal layer (simple linear regression, *R*^2^ = 0.0 for MEM, Sham, *R*^2^ = 0.006 for MPEP, Sham), while it did not affect interlayer spatial-functional correlation (Table S8). As we observed for neuronal and multilayer topology, pre-treatment of injured dishes with MPEP attenuated the full effect of injury for the neuronal layer (simple linear regression, *R*^2^ = 0.006 for MEM, Inj, *R*^2^ = 0.001 for MPEP, Inj) and interlayer connections (simple linear regression, *R*^2^ = 0.007 for MEM, Inj, *R*^2^ = 0.0 for MPEP, Inj) Together, these results indicate that injury triggers a process of forming more local networks, implying a loss in the coordination of signaling across the networks, that is influenced by mGluR_5_ signaling.

### Modularity analysis reveals community association of neurons and astrocytes is not determined by spatial proximity at the mesoscale

In addition to local and global topological statistics, we sought to understand the community structure of multilayer networks, which can be used to study dynamics in a subset of a larger network. Further, we asked whether community structure depends on the physical distance between nodes. To do so, we compared the modularity predicted by functional and spatial connectivity. The modularity of both functional and spatial graphs was determined by maximizing the quality function (Eq. 12), constructed with a Newman-Girvan null model (Newman and Girvan (2004)) (Fig. S13), using a Louvain-like locally greedy algorithm implemented in MATLAB (Jutla, Jeub, and Mucha (2011)). The spatial tuning parameter was adjusted to minimize the difference between the number of communities in the spatial networks and the number of communities in the functional networks. To compare the modularity predicted by spatial and functional graphs, we calculated the Adjusted Rand Index (*ARI*, Rand (1971)) between the partitions of spatial and functional networks into communities (Eq. 16). The Rand Index is equal to one when two community partitions are exactly equal, and zero when there is no overlap. The *ARI* adjusts for the probability of two cells being assigned to the same module by chance, and can take on negative values.

Despite a small but significant correlation between functional and spatial connectivity, we found that community structure differed between functional and spatial networks for neuron-neuron networks (Fig. S7E-F), with the *ARI* between functional and spatial partitions being no different from zero (neuron networks, mean = 0.01311, two-sided one-sample *t*-test, *t* = 1.502, *df* = 35, *p* = 0.1420. The mean *ARI* between spatial and functional multilayer communities was near but significantly greater than zero (mean = 0.05364, one sample *t*-test, *t* = 4.395, *df* = 22, *p* = 0.0002). This disagreement between spatial and functional partitioning was not driven by differences in community size, as *γ*, the spatial tuning parameter, was adjusted to minimize this difference (Fig. 4H, paired *t*-test, *t* = 0.1645, *df* = 22, *p* = 0.8709; see also SI Sec. F). This finding suggests that the functional connectivity of groups of neurons and astrocytes is largely independent of distance at this scale *in vitro*. Furthermore, the choice to use the multilayer modularity approach allowed us to calculate cell type specific participation across modules (Fig. 4I). Relative to astrocyte segments, neurons participated in a greater fraction of functional (Tukey’s multiple comparisons test, *q* = 8.549, *df* = 88, *p <* 0.0001) and spatial (Tukey’s multiple comparisons test, *q* = 11.28, *df* = 88, *p <* 0.0001). This difference in module participation reflects the dominance of neurons in quantity, as sub-sampled networks with an equal number of neurons and astrocytes had near equal spatial and functional participation of both cell types (SI Sec. E, Fig. S6E).

Because astrocytes were subdivided into smaller functional units of microdomains, their functional modularity could be directly compared to their morphological (actual) modularity, which serves to group segments of the same cell (Fig. 5A-B). This approach enabled us to test whether microdomains of the same astrocyte are functionally independent at the mesoscale. To test whether functional and spatial graphs of astrocyte segments matched actual cell morphology, we computed the *ARI* between functional, spatial, and actual astrocyte communities. Astrocyte microdomains were manually grouped into whole cells based on maximum fluorescence projection images (Fig. S3). Community partitions between the three graph types (functional, spatial, and actual) were different, with *ARI <* 0.4 for all (Fig. 5C). Interestingly, the *ARI* between cellular and functional communities was largest, significantly larger than cellular versus spatial (Tukey’s multiple comparisons test, *q* = 3.369, *df* = 132, *p* = 0.0486) and spatial versus functional (Tukey’s multiple comparisons test, *q* = 4.531, *df* = 132, *p* = 0.0048) community partitions generated from multilayer networks. Differences in community membership were not predominantly attributable to changes in community size or community number between graph types, as *γ* was tuned to minimize differences in mean community size of the three partitions at baseline (one-way ANOVA, *F* (2,66) = 0.08331, *p* = 0.9202). Differences in actual, spatial, and functional modularity suggest that astrocyte segments behave relatively independently from the rest of the cell, and tend to co-function with neurons.

**Figure 5.**
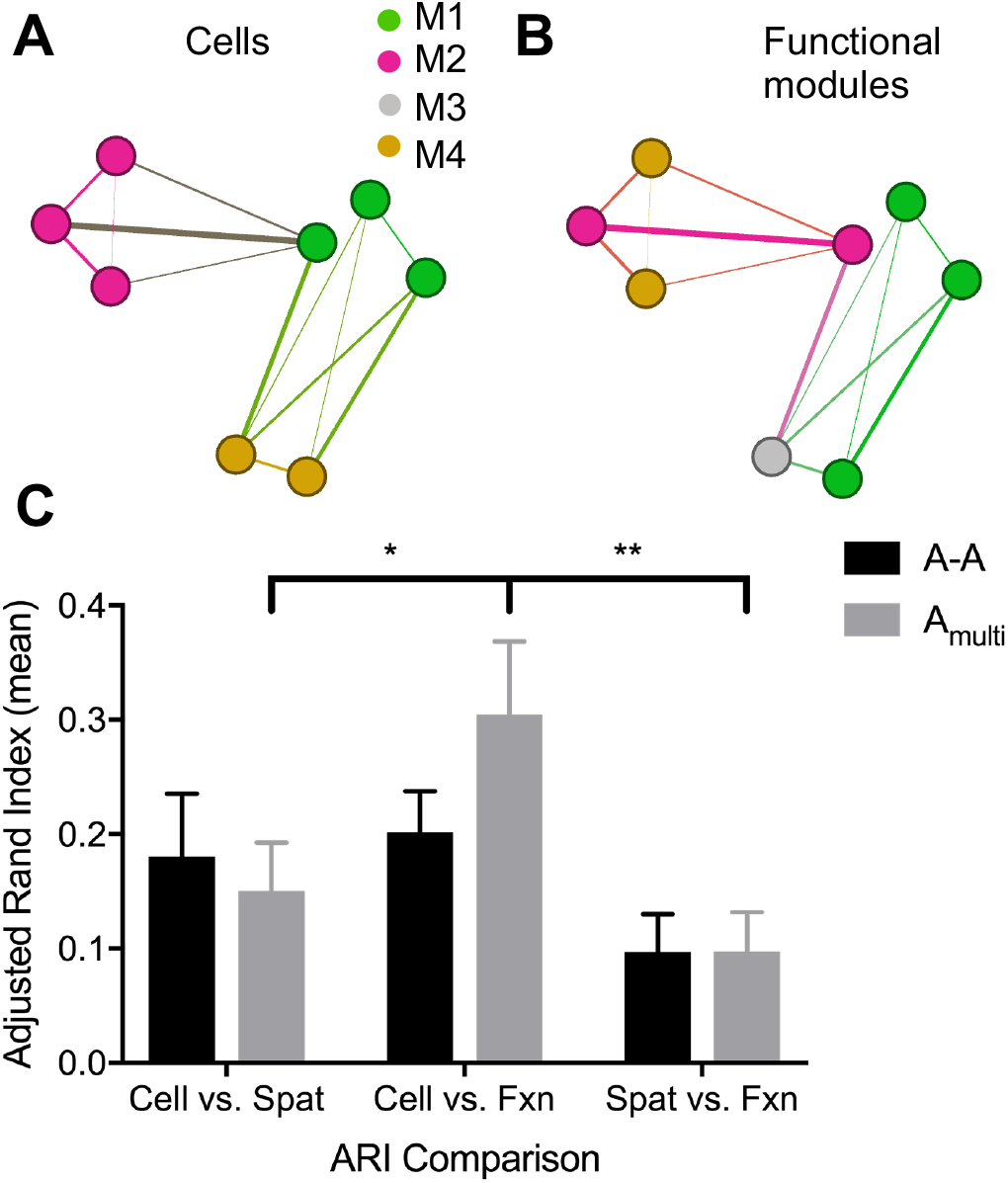
Functional and spatial communities of astrocyte segments are independent of cell membership, but less so in multilayer networks. **A**. A representative astrocyte network with the indicated community structure based on morphology of segments. Nodes (astrocyte segments) of the same color belong to the same cell. **B**. The network in panel **A** with modules detected based on functional connectivity. Nodes (astrocyte segments) of the same color belong to the same functional module, independent of cell membership. **C**. Mean Adjusted Rand Index (*ARI*) for morphologically connected astrocyte segment communities (cell) versus spatially-connected communities, morphologically-connected versus functionally-connected segment communities, and functionally-connected versus spatially-connected communities. Morphological versus spatial *ARI* is larger than morphological versus functional, and also than functional versus spatial *ARI*s in both independent astrocyte (black) and multilayer (gray) astrocyte modules (two-way ANOVA comparing the effect of module type and single- or multilayer network on *ARI*: *F* (1,132) = 7.675, *p* = 0.0062). The *ARI* is always larger for astrocyte communities generated using a multilayer network, though the improvement in *ARI* is not statistically significant (two-way ANOVA comparing the effect of single versus multilayer network type on *ARI*: *F* (2,132) = 2.911, *p* = 0.0579). Error bars indicate standard error of the mean (SEM) and asterisks indicate statistical significance (*p *≤* 0.05, **p *≤* 0.01, ***p *≤* 0.001, ****p *≤* 0.0001).

When multi- and single-layer astrocyte *ARI*’s were grouped, differences between module types were significant (two-way ANOVA comparing the effect of module type and single or multilayer network on *ARI*: *F* (1,132) = 7.675, *p* = 0.0062). This finding highlights the functional independence of astrocyte segments of the same cell in both their own layer and in the multilayer network. Further, *ARI* between all module types was improved when astrocyte communities were assigned from multilayer graphs, although this improvement was not statistically significant (two-way ANOVA comparing the effect of single versus multilayer network type on *ARI, F* (2,132) = 5.840, *p* = 0.0037). That is, incorporating information about neural activity improved the agreement between morphologically and functionally determined astrocyte segment modules. One interpretation of this finding is that functional connectivity between astrocyte segments and neurons is more influenced by spatial proximity than is functional connectivity between astrocyte segments. Furthermore, this improvement in morphology-function agreement in multilayer astrocyte communities, which include neurons, suggests that the two populations are not functionally independent. If they were independent, we would expect the community structure of astrocyte segments to remain unchanged after multilayer modularity analysis.

### mGluR_5_ localizes to neurons and astrocytes in culture

To verify the presence and assess the relative abundance of mGluR_5_ on neurons and astrocytes in our cultures, we stained for mGluR_5_; microtubule associated protein 2 (MAP2), a neuronal marker; glial fibrillary acidic protein (GFAP), an astrocytic marker; and nuclei (Hoechst stain), for counting purposes. Six dishes of näive cells were fixed at DIV 10, the same age at which experimental cells were treated and imaged, stained using immunofluorescent antibodies (SI Sec. G), and imaged in three fields of view.

MAP2 and mGluR_5_ co-localized in 99% of neurons (*n* = 6 dishes, *n* = 3 fields of view each, *n* = 525 neurons total), verifying expression of the receptor on neurons. GFAP and mGluR_5_ co-localized in 76% of astrocytes (*n* = 6 dishes, *n* = 3 fields of view each, *n* = 119 astrocytes total), confirming the expression of the receptor in astrocytes. However, anti-mGluR_5_ intensity was visibly lower in astrocytes than in neurons (Fig. S8), suggesting that neurons in our cultures express more mGluR_5_ than astrocytes. A notable limitation of GFAP as an astrocyte marker is that it does not stain fine astrocyte processes or microdomains (Shigetomi et al. (2013)), where mGluR_5_ is more likely to be located (Panatier and Robitaille (2016); Panatier et al. (2011)). Thus, our stain may not have captured the full extent of astrocytic mGluR_5_ expression.

## DISCUSSION

Using an *in vitro* model of traumatic brain injury, this study examined the functional response of cultured neuron-astrocyte populations to disruption of mGluR_5_-mediated signaling and mechanical injury. mGluR_5_ inhibition decreased neuronal, but not astrocytic, activity following injury, but did not alter higher-order topological properties of multilayer networks on its own. In contrast, mechanical injury to neurons increased the global efficiency and clustering of the neuronal layer and clustering of the entire multilayer network, but did not affect the topology of the astrocyte layer. Pre-treatment of injured cultures with an mGluR_5_ inhibitor attenuated the effect of injury on neuronal and multilayer topology. In order to determine whether network connectivity between cells was primarily determined by spatial proximity at both the individual node and community scales, we compared the functional connectivity and community structure of networks generated using calcium activity and spatial location. This analysis revealed that functional connectivity is largely independent of spatial proximity in small-scale *in vitro* neuron-astrocyte networks. However, for both layers of the multilayer network and for interlayer connections, mechanical injury increased the extent to which functional connectivity correlated with spatial proximity. This increased spatial dependency was attenuated by pre-treatment with an mGluR_5_ inhibitor for the neuronal layer and interlayer connections. Finally, we observed that segments of the same astrocyte often interact with separate functionally connected communities of neurons. Our results suggest that while astrocytes and neurons are functionally connected, mGluR_5_ is not the primary receptor mediating neuron-astrocyte communication in our culture system. In this discussion, we explore potential explanations for our findings and their significance in the context of glial biology and traumatic brain injury.

### Disrupted neuronal functional connectivity following mechanical injury, modulated by mGluR_5_ blockade

Previous studies using other in vitro models have shown that mechanical injury to neurons reduces functional connectivity (W. H. Kang et al. (2015); Myer et al. (2006); Patel et al. (2012)). In line with these studies, our results show that single-cell tap injury to multiple neurons alters network topology in favor of increased clustering and larger global efficiency in the neuronal layer. Certainly, the precise removal of individual neurons from a network would decrease the synaptic inputs to neighboring neurons, leading to an acute reduction in the activity of the network. However, this could be followed by a robust homeostatic re-scaling of network (Turrigiano and Nelson (2004)) which could subsequently alter the functional connectivity of the remaining neurons. Alternatively, randomly removing inhibitory neurons within the network, with their more restricted range of spatial connections, may also reorganize the activity to establish bursts of activity within smaller subnetworks, resulting in a network with an increased number of hubs, an increased efficiency, and an increase in clustering. Regardless of its exact origin, our results demonstrate the intricate interruption in balance across neuronal, but not astrocytic, layers that occurs from microtrauma to neurons.

Our observation that injured networks pre-treated with MPEP did not exhibit the changes in topology observed for injured networks suggests that the mGluR_5_ receptor facilitates the neuronal response to injury. Specifically, the increased mean nodal strength, clustering, and global efficiency of neuronal layers following injury is likely to be facilitated, at least in part, by glutamatergic signaling via mGluR_5_. Because we do not observe these changes in the astrocyte layer or in inter-layer strength, we speculate that neuronal mGluR_5_ is likely responsible. This finding is in agreement with previous work, which has found that long-term potentiation (LTP) at thalamic input synapses was impaired following bath application of MPEP in vitro (Rodrigues et al., 2002). In *in vivo* follow-up, administration of MPEP dose-dependently impaired the acquisition of auditory and contextual fear learning, implicating mGluR_5_ in neuronal plasticity, and highlighting the functional importance of this receptor. In another study, mGluR_5_-deficient mutant mice showed a complete loss of the NMDA-receptor mediated component of LTP in hippocampal CA1 neurons (Jia et al. (1998)). From these and our results, we infer that mGluR_5_ plays an important role in establishing and maintaining the functional connectivity of neural circuits.

While injury did not alter astrocyte population activity, it did change the topology of multilayer networks in a manner that was not observed completely in neuron or astrocyte populations alone. Specifically, injury increased mean nodal strength, clustering, and global efficiency in the neuronal layer, but did not significantly affect mean strength of multilayer, interlayer, or astrocyte layer nodes. However, injury did increase the clustering, but not global efficiency, of multilayer networks. Taken together, these results suggest that multilayer changes in clustering coefficient are mostly mediated by the neuronal layer. Conceptually, increased mean nodal strength in the neuronal layer following injury may result from local changes in synaptic strength from plasticity, or may also result from NMDA receptor subtypes influencing the remodeling of synaptic strength. On its own, spike timing dependent plasticity can compensate quickly for reductions in activity that occur when a fraction of the neuronal population is inactivated, leading to a recovery in connectivity within the microcircuit and an increase in remaining synaptic strength (Gabrieli, Schumm, Vigilante, Parvesse, and Meaney (2020); Schumm, Gabrieli, and Meaney (2020)), in turn increasing the functional connectivity in the network. Our model of mechanical trauma may lead to local increases in functional connectivity within neurons containing a large fraction of NMDAR receptors containing the GluN_2_A subunit (Patel et al. (2014)). Injury also increased the clustering coefficient, reflective of closure in local neighborhoods in the multilayer network. These topological changes may result from functionally eliminating nodes through which long-distance paths pass, forcing more of the remaining healthy neuronal nodes to adopt hub roles, thereby increasing local clustering. In this light, the elimination of nodes appears to promote a rerouting information through other healthy nodes to indirectly trigger a ‘pathological plasticity’, where increases in functional connectivity are a necessary step in rebuilding the network. The interaction of injury and drug did not have this effect on clustering, suggesting that mGluR_5_ inhibition prevents the impact of injury on topology. This protective effect of mGluR_5_ inhibition following injury was observed in previous studies *in vitro* and *in vivo* (Lea IV, Custer, Vicini, and Faden (2002); Movsesyan et al. (2001)), and has been attributed to a reduction in NMDAR activity following mGluR_5_ inactivation.

### Unchanged neuron-astrocyte connectivity following mGluR_5_ inhibition

While molecular and cellular changes following direct injury to astrocytes is well-documented (Charles et al. (1991); Ellis et al. (1995); Rzigalinski et al. (1998)), the functional contribution of astrocytes to neural population activity in response to targeted neuronal injury is unknown. Due to the expression of mGluR_5_ on neurons and astrocytes, we were unable to target astrocyte-specific mGluR_5_ pharmacologically. Thus, we could not determine whether changes in neuronal network density were attributable to changes in gliotransmission. Contrary to our initial hypothesis, astrocyte activity and functional connectivity did not decrease following injury or mGluR_5_ inhibition. Because we did not directly injure astrocytes, their lack of direct response to injury after one hour is perhaps expected. However, the lack of astrocyte response to mGluR_5_ blockade is more difficult to interpret. It could be that astrocytes are able to compensate for impaired glutamatergic signaling through other modes of intra-astrocyte communication, including gap junctions and other biochemical receptor pathways. Alternatively, we may have examined astrocyte activity at too coarse of a spatial scale. Astrocyte mGluRs are localized in microdomains, which we were unable to visualize under cytosolic GCaMP6f or GFAP staining. Thus, we likely did not observe the full extent of astrocyte activity and its dynamics following injury and mGluR_5_ inhibition. Improper spatial resolution might also explain the low expression of mGluR_5_ on astrocytes relative to neurons observed in stained cultures. Additionally, previous work indicating a critical role for mGluR_5_-mediated neuron-astrocyte communication was primarily conducted in slices and in vivo, rather than in dissociated cultures. Thus, it is possible that different gliotransmission pathways, perhaps purinergic (Panatier and Robitaille (2016)), are relatively more important in our cultures, which differ in age, health, cellular composition, and genetic variation from astrocytes in slices and *in vivo*.

### Independence of spatially and functionally predicted community structure in small-scale neuron-astrocyte networks

Although significant progress has been made towards understanding spatially embedded networks and community structure *in vivo* (Bassett and Stiso (2018); Betzel and Bassett (2017)), functionally and structurally informed modularity of *in vitro* neuronal networks has not been explored. Because cultured cells exist in a constrained 2-dimensional environment, devoid of connective tissue and vasculature, the spatial organization of *in vitro* networks is likely to be different from that of *in vivo* networks. Furthermore, we examined a small sub-section of a larger network in this study. In this context, it is logical that the functional and spatial community structure of neuron-astrocyte networks are mostly independent. Neurons and astrocytes can connect with one another over distances an order of magnitude larger than the approximately 1mm field of view imaged in this study. Thus, functional connectivity can be established largely independent of spatial proximity in healthy networks. It could be that spatial proximity strongly influences functional modularity at and above a certain scale, or when the network is under stress. Comparing spatial and functional network modularity in astrocyte segments facilitated more general characterization of astrocyte calcium signaling. Astrocytic calcium events are heterogeneous in temporal and spatial profile, ranging from fast, locally restricted events to slower, cell-wide waves (Bazargani and Attwell (2016); Stobart et al. (2018); Y. Wang et al. (2018)), with these two types of events often occurring independently (Di Castro et al. (2011); Panatier et al. (2011)). There is limited understanding of how Ca^2+^ fluctuations in microdomains relate to cell-wide, or population-wide, astrocyte calcium events (Nimmerjahn and Bergles (2015)). Analyzing astrocyte segments as nodes in a spatially embedded network enabled us to elucidate the extent to which segments of the same astrocyte are functionally interdependent. If segments of the same astrocyte participate in different functional modules, there is likely a functional advantage to maintaining strong inter-astrocyte and astrocyte-neuron connectivity above intra-astrocyte connectivity. This microdomain independence allows a given astrocyte to sense and integrate activity from a diverse set of neurons over time and space (Araque et al. (2014); Fields, Woo, and Basser (2015)). Multilayer astrocyte modularity recapitulated this mesoscale structure-function independence, but to a lesser degree.

### Methodological considerations

A major limitation of this study is the extreme sparseness of active astrocyte segments relative to neurons in our cell cultures. Neuronal dominance of multilayer network dynamics likely skewed multilayer topology towards the neuronal layer. The low level of activity in astrocytes could be due to our examination of astrocyte activity at specific spatial and temporal scales or to the health of the cells in culture. Prior work indicating critical roles for neuron-astrocyte communication in plasticity, synchronization, and behavior, were primarily conducted in slices and *in vivo*, and many involved direct stimulation of neurons or promoted arousal to evoke an astrocyte response (Srinivasan et al. (2015); X. Wang et al. (2006)). In contrast, we examined relatively asynchronous spontaneous neural activity in dissociated cultures. The astrocytes in our model system may not have developed the complete range of neuromodulatory activities performed by those *in vivo*.

The interpretation of our findings depends on relative mGluR_5_ expression and signaling importance in the neurons and astrocytes in our cultures. The presence and importance of mGluR_5_ on neurons and in their signaling is well-documented (Ballester-Rosado et al. (2010); Gass and Olive (2009); Shigemoto et al. (1993)) and has been confirmed with immunofluorescent staining of mGluR_5_ in our *in vitro* preparations. Indeed, the decrease in neuronal network density following MPEP administration is likely attributable to blockade of neuron-neuron communication. This is an inherent limitation in conventional pharmacological studies of gliotransmission. To isolate the causal effect of astrocyte-specific signaling on neural circuit activity, future experiments could use more sophisticated tools such as designer receptors exclusively activated by designer drugs (DREADDs) (Scofield et al. (2015)), astrocyte-specific G-protein coupled receptor opsins (Mederos et al. (2019)), or genetic knockdown (Shigetomi, Tong, Kwan, Corey, and Khakh (2012)).

### Future directions: in vivo network models of neuron-astrocyte signaling and expanded models of neuron-glia networks

The aforementioned differences in health and activity between astrocytes *in vitro* and *in vivo* motivate future work comparing the topology and response behavior of neuron-astrocyte networks in cell culture and animal models. Assuming that the number of active astrocyte microdomains *in vivo* is greater than in culture, we speculate that there would be significant changes in neuron-astrocyte network community structure, as astrocytic participation in spatial and functional models would increase. Similarly, we speculate that there is increased density and strength of astrocyte-astrocyte and neuron-astrocyte connections *in vivo*. Given our findings that betweenness centrality, global efficiency, and clustering are mostly influenced by density and strength, these topological properties would change accordingly in *in vivo* networks. Furthermore, with greater neuron-astrocyte connectivity *in vivo*, neuronal manipulations, such as injury, would induce a greater change in multilayer and astrocyte layer network topology. Additionally, we propose that the topology of neuron-astrocyte networks changes from its spontaneous state during periods of intense stimulation, given that astrocytes exhibit unique responses to stimulation and during behavior (Halassa et al. (2009); Lee et al. (2014); Perea, Yang, Boyden, and Sur (2014); Schummers, Yu, and Sur (2008); Srinivasan et al. (2015); X. Wang et al. (2006)). For example, there may be increased alignment between morphological and functional astrocyte communities during stimulation, as repeated neuronal firing triggers global astrocyte Ca^2+^ events (Khakh and Sofroniew (2015)). Indeed, astrocytes *in vivo*, which occupy non-overlapping territories, have been hypothesized to form functional “islands” of neuronal synapses (Halassa, Fellin, Takano, Dong, and Haydon (2007)). It is difficult to speculate how our findings related to mGluR_5_ inhibition will translate *in vivo*. The relevance of mGluR_5_-mediated gliotransmission in adult rats is debated, as there is evidence of decreased mGluR_5_ expression after 7-21 days postnatally (Cai, Schools, and Kimelberg (2000); Morel, Higashimori, Tolman, and Yang (2014); Sun et al. (2013)).

The field of network neuroscience was largely built upon observations of structural connectivity at the whole brain or macro-scale represented as graphs (Bassett and Bullmore (2017); Bassett et al. (2018); Liao, Vasilakos, and He (2017)). Combined with the ever-increasing focus on data science, the field has left first-principles descriptions of functional connectivity at the elementary (single-neuron) level relatively underexplored (Bassett et al. (2018)) Indeed, there have been few functionally-derived network models of neuronal microcircuits (Bettencourt, Stephens, Ham, and Gross (2007); Bonifazi et al. (2009); W. H. Kang et al. (2015); Patel et al. (2012); Schroeter, Charlesworth, Kitzbichler, Paulsen, and Bullmore (2015); Srinivas, Jain, Saurav, and Sikdar (2007)). Our work aids in addressing this understudied area by characterizing the topology, and changes therein, of functional connections between elementary-scale mixed cortical cultures. Further, to our knowledge, the field of network neuroscience is devoid of any models describing or predicting connectivity between neurons and glia, although there are some computational neuroscience models that explore the interactions of these two cell types. Network modeling of neuron-glia interactions is an area ripe for future study, and is particularly well suited for characterization and prediction using multilayer models.

The field of network neuroscience has used multilayer networks to model brain network development over time, to compare healthy and diseased brains, to predict structure–function relationships, and to link multi-scale and multi-modal data (Bassett et al. (2011); Betzel and Bassett (2017); Betzel et al. (2018); Muldoon and Bassett (2016); Pedersen, Zalesky, Omidvarnia, and Jackson (2018)). Despite the increasing popularity of multilayer networks and the existence of multiple neural cell types, there has been limited use of the multilayer framework to study interactions between two or more cell populations. Here, we demonstrate the validity of a multilayer network model to characterize the connectivity of neuron-astrocyte populations.

## Conclusion

Our results indicate that mGluR_5_ is not essential for maintaining spontaneous astrocyte-astrocyte or neuron-astrocyte functional connectivity in non-injured cultures. However, mGluR_5_ blockade partially restored baseline topology of multilayer networks following traumatic injury, underscoring the potential therapeutic benefit of modulating glutamatergic tone. More broadly, we have proposed and demonstrated the utility of a multilayer network model of *in vitro* neuron-astrocyte populations in interpreting the topological impacts of various experimental manipulations. Our findings support the notion that neurons and astrocytes are not functionally distinct populations, but rather display an interdependent topology that reveals network responses not observed in either population alone. Future work could aim to isolate the astrocytic contribution to static and dynamic network topology using more targeted experimental manipulations, and could study healthy neuron-astrocyte populations *in vivo*, where astrocytes may be more abundant and active.

## MATERIALS AND METHODS

### Experimental protocol

All animal procedures were approved by the University of Pennsylvania Institutional Animal Care and User Committee. Timed pregnant Sprague-Dawley rats were anesthetized with 5% CO_2_ and sacrificed via cervical dislocation. Embryos at day E18 were surgically removed and their neocortical tissue dissected and dissociated for 15min at 37°C in trypsin and DNAse. Following trituration and filtration through Nitex mesh, cells were re-suspended in minimum essential media (MEM) with Earl’s salts and GlutaMAX supplemented with 0.6% D-glucose, 1% Pen-Strep, and 10% Horse Serum. Cells were plated on glass-bottomed Matteck dishes coated in poly-D-lysine and laminin at a density of 300,000 cells/mL. Cultures were grown in a humidified incubator at 37°C and 5% CO_2_, maintained in serum-free MEM and 0.2% GlutaMAX (Gibco), and fed twice weekly with 1X NEUROBASAL medium (ThermoFisher).

To visualize the calcium activity of astrocytes in addition to neurons, cells were transduced using Adeno-associated virus Serotype 1 (AAV1) carrying the gene for GCaMP6f under a CAG promoter, which is general to mammalian cells (AAV1.CAG.Flex.GCaMP6f.WPRE.SV40, Addgene), at DIV 3. All manipulations and imaging were performed at DIV 10. Dishes were randomly chosen to be treated with 1uM 2-Methyl-6-(phenylethynyl)pyridine (MPEP), a potent mGluR_5_ inhibitor (for 19 dishes), or MEM only (n = 36 dishes total, Fig. S10). Calcium images were obtained at 488nm excitation, 50ms exposure on a Nikon Eclipse TE2000 confocal microscope. Eight to nine dishes in each treatment group were mechanically injured by briefly deforming the cell surface of 12-15 neurons with a vertical movement of a pulled glass micropipette tip (Charles et al. (1991)). The micropipette was controlled manually with a microcontroller (Eppendorf), directly perturbing the cell with the pipette tip. Care was taken to avoid rupturing the cell membrane, which would have caused immediate cell death. This model enabled precise, controlled injury of a similar severity across dishes.

### Image processing

Images were manually segmented to identify neurons and astrocyte microdomains as regions of interest (ROIs, Fig. S9, Fig. S3). Neuronal and astrocytic fluorescence traces were extracted from each ROI as pixel intensity over time (Fig. S9). Neuronal and astrocytic events were identified automatically, using a custom-built MATLAB template-matching algorithm (Patel et al. (2015)), based on the trace’s correlation with a library of true neuron and astrocyte calcium waveforms (Fig. S11, Fig. S12). Automatically identified events were manually confirmed based on calcium traces or eliminated following event detection. Because astrocytic activity was mostly localized to processes that often fluoresced independently, an observation that has been confirmed in previous studies (Fields et al. (2015); Srinivasan et al. (2015)), astrocyte segments were analyzed as separate entities. Cells predicted to be astrocytes based on morphology were confirmed based on their functional response to 100uM NMDA administered with 1uM of glycine co-agonist (SI Sec. B). Only ROIs that had one or more confirmed calcium events in one or more recordings were kept for further analysis. This pruning step eliminated a significant number of nonactive astrocyte segments, reducing the number of dishes with two or more active astrocyte segments to 23 from 36.

### Generation of adjacency matrices

We based functional connectivity on filtered, scaled calcium fluorescence. For a pair of ROIs, either neuronal or from an astrocyte microdomain, we calculated the Pearson’s correlation coefficient between the filtered, scaled calcium fluorescence trace and 100 surrogates with similar properties to the original traces. Specifically, the surrogate time series had the same marginal distribution and power spectrum as the original, and generated using an amplitude adjusted Fourier transform (AAFT) algorithm (Kugiumtzis and Tsimpiris (2010)). Only pairs of ROIs that had a phase difference significantly larger or smaller than the surrogates, as determined by a *z-*test with *p <* 0.05, were linked with an edge having a weight equal to the correlation between the two traces (SI Sec. C). Multilayer adjacency matrices were organized as shown in Fig. 2, with neuron-neuron, neuron-astrocyte, and astrocyte-astrocyte adjacency matrices concatenated. Spatial adjacency matrices were generated by measuring the radial distance between the center of each region of interest (ROI) of each node pair, (*x*1, *y*1) and (*x*2, *y*2), as √(|(*x*2−*x*1)|^2^+|(*y*2−*y*1)|^2^). The distance was normalized to the maximum possible distance in the field of view, and subtracted from one so that the pairs of nodes with the shortest radial distance were linked by edges with the highest weights. Null networks were generated by randomizing adjacency matrices while preserving degree distribution and density using the *randmio* function in the BCT (Maslov and Sneppen (2002)). Edges in networks with more than three edges were rewired 100 times for neuron networks, 10 times for astrocyte networks due to their smaller size, and 100 times for multilayer networks.

### Calculation of network statistics

Density, strength, clustering coefficient (*C*), betweenness centrality (*B*), and global efficiency (*E*) were calculated for neuron, astrocyte, and multilayer networks using weighted adjacency matrices. Equations are provided in the SI Sec. D. In most cases, MATLAB scripts from the BCT were used for calculations (Rubinov and Sporns (2010)). The form of the multilayer adjacency matrices lends itself to straightforward calculation of network statistics in the same way as for single-layer networks. To compare the topology of the two layers and intra- vs. interlayer connections, we calculated mean nodal strength of the entire network, mean nodal strength of each individual layer, and mean strength of interlayer connections. Interlayer strength and density were calculated by summing the weights, or number of nonzero values, of the edges between neurons and astrocytes, and was divided by (*N*_*n*_ + *N*_*a*_)*/*2, where *N*_*n*_ is the number of neurons and *N*_*a*_ is the number of astrocytes, to normalize for network size. More global network statistics, such as betweenness centrality, global efficiency, and clustering coefficient, cannot be separated into intra- and interlayer components. These metrics were calculated for the entire multilayer network and also for individual layers. To estimate the effect of treatment group assignment on these graph statistics (*B, C*, and *E*), we ran a generalized linear regression Nelder and Wedderburn (1972) using the GLM module class in Python’s *statsmodels* module Seabold and Perktold (2010). The link function was the identity and error distribution was assumed to be normal. Outcome variables *B, C*, and *E* were regressed against categorical variables indicating treatment group (MEM, No Injury (Sham); MPEP, No Injury (Sham); MEM, Injury; or MPEP, Injury), as well as mean nodal strength, to control for its effects on topological statistics. Regression coefficient estimates and *p*-values are reported above.

### Community detection in spatial and functional graphs

To better understand how individual astrocyte segments and neurons give rise to collective network structure, we analyzed the multilayer network’s mesoscale community structure (Betzel and Bassett (2017); M. A. Porter, Onnela, and Mucha (2009), Fig. S13). Modules are communities of nodes that are functionally or spatially more interconnected to each other than to other nodes in the network. Modules were generated by maximizing the modularity quality function using both spatial and functional adjacency matrices for single-layer neuron and astrocyte networks, and for multilayer networks. The parameter *γ* is a resolution parameter that governs the size and number of detected communities. For both single-layer and multilayer networks, *γ* was tuned separately and manually for spatially and functionally generated graphs to minimize the difference between the number of communities detected for spatial and functional networks (SI Sec. F). Quality was maximized by placing nodes into communities such that the quality *Q* is as large as possible, using an iterative Louvain-like community detection algorithm implemented in MATLAB Jutla et al. (2011). A Girvan-Newman Newman and Girvan (2004) null model was used for both functional and spatial graphs and implemented in a custom MATLAB script (SI Sec. F). Because the modularity landscape is rough Good, De Montjoye, and Clauset (2010), the heuristic can detect a different partition each time it is applied. Thus, here we assign a node’s module to be the mode over 50 maximizations. Nevertheless, module assignment still varies slightly between batches of 50 trials, a property that more generally has been demonstrated for small networks Good et al. (2010). In addition to the value of *Q*, the Louvain algorithm outputs the partition *g*, a vector containing the community number of each node.

We compared the similarity of the partitions predicted by functional and structural graphs by calculating their Adjusted Rand Index (*ARI*) Hubert and Arabie (1985); Rand (1971) using a MATLAB algorithm “RANDINDEX” (2000). The Rand Index represents the probability that the two partitions, functional and spatial, will agree on the community of a randomly chosen node. A Rand Index of zero indicates that the two partition vectors do not have any groupings in common, while a value of one indicates complete agreement. The *ARI* corrects for over-inflation due to chance assignment (SI Sec. F). Module participation for each cell type was calculated as the fraction of all modules that contain at least one of that cell type.

### Statistical testing

Two-sided one-sample *t*-tests were conducted to determine the statistical significance of the difference between actual and null network statistics, as well as the statistical significance of the difference of the mean *ARI* from zero for neuron layer (*n* = 36 dishes), astrocyte layer (*n* = 23 dishes) and multilayer (*n* = 27 dishes) networks. Two-sided paired *t*-tests were conducted to determine whether there existed a statistically significant difference between the size of functionally and spatially determined modules for neuron (*n* = 36 dishes), astrocyte (*n* = 23 dishes), and multilayer (*n* = 27 dishes) networks. A two-sided unpaired *t*-test was used to test for a statistically significant difference between the size of multilayer network functional communities in MPEP- and MEM-treated dishes. One-way analysis of variance (ANOVA) was used to test for a significant difference between means of a single measure between multiple groups. Two-way ANOVA was used to test for a significant difference between means of multiple measures between multiple groups. The Holm-Sidak method was used to correct for multiple comparisons when making post-ANOVA comparisons between the means of selected pairs of groups.

The Tukey method was used to correct for multiple comparisons when making post-ANOVA comparisons between every mean and every other mean. Two-sided *t*-tests were used to test for statistical significance of all regression coefficients.

## ACKNOWLEDGMENTS

We thank S. Schumm, K. Wiley, and P. Srivastava for helpful comments on an earlier version of this manuscript; A. Nam for primary cell culture; and O. Teter for collaborative brainstorming.

This work was supported by a Paul G. Allen Family Foundation grant to DFM and DSB. DFM would like to acknowledge the support of the NINDS (1RO1-NS 088276). DSB would also like to acknowledge support from the John D. and Catherine T. MacArthur Foundation, the Alfred P. Sloan Foundation, the ISM Foundation, the Army Research Laboratory (W911NF-10-2-0022), the Army Research Office (Bassett-W911NF-14-1-0679, Grafton-W911NF-16-1-0474, DCIST-W911NF-17-2-0181), the Office of Naval Research, the National Institute of Mental Health (2-R01-DC-009209-11, R01-MH112847, R01-MH107235, R21-M MH-106799), the National Institute of Child Health and Human Development (1R01HD086888-01), National Institute of Neurological Disorders and Stroke (R01 NS099348), and the National Science Foundation (BCS-1441502, BCS-1430087, NSF PHY-1554488 and BCS-1631550)). The content is solely the responsibility of the authors and does not necessarily represent the official views of any of the funding agencies.

## SUPPORTING INFORMATION

### Supplemental Text

Fig. S1. Calcium activity and size of neuron-astrocyte cultures.

Table S1. Results of Tukey’s multiple comparisons test following a 2-way ANOVA on the effect of time and group assignment on neuronal and astrocytic event rate at the final experimental time point (one hour post-injury).

Fig. S2. Dependence of observed and randomized network topology on edge density.

Table S2. Results of linear regressions of *B, C*, and *E* on network density *κ* for observed (obs) and randomized (rand) networks at the final experimental time point.

Fig. S3. Manual grouping of astrocyte microdomains.

Fig. S4. Effect of exogenous manipulations on neuron network edge density and nodal strength.

Fig. S5. Effect of exogenous manipulations on astrocyte network edge density and nodal strength.

Table S3. Results of generalized linear regression to predict the effect of group assignment on mean clustering coefficient, mean normalized betweenness centrality, and global efficiency for neuron networks at the final experimental timepoint.

Table S4. Results of generalized linear regression to predict the effect of group assignment on mean clustering coefficient, mean normalized betweenness centrality, and global efficiency for astrocyte networks at the final experimental timepoint.

Table S5. Results of generalized linear regression to predict the effect of group assignment on mean clustering coefficient, mean normalized betweenness centrality, and global efficiency for multilayer networks at the final experimental timepoint.

Fig. S6. Characterization of sub-sampled multilayer neuron-astrocyte functional and spatial network topology.

Table S6. Results of simple linear regression of *B, C*, and *E* on network density *κ* for sub-sampled multilayer networks at the final experimental time point.

Table S7. Results of generalized linear regression to predict the effect of group assignment on mean clustering coefficient, mean normalized betweenness centrality, and global efficiency (E) for sub-sampled multilayer networks at the final experimental timepoint.

Fig. S7. Community structure of *in vitro* neuronal networks.

Table S8. Results of linear regressions to predict functional adjacency from spatial adjacency for each experimental group at the final experimental time point.

Figure S8. Immunofluorescent staining confirms expression of mGluR_5_ in neurons and astrocytes.

Fig. S9. *In vitro* calcium image acquisition and processing of neuron-astrocyte networks.

Fig. S10. Experimental design and treatment protocol.

Fig. S11. Spike inference from neuronal calcium activity.

Fig. S12. Event inference from astrocyte calcium activity.

Fig. S13. Community detection in a multilayer functional network.

## DATA AND CODE AVAILABILITY

Data is available from the corresponding author upon reasonable request. Custom MATLAB scripts are available for download at https://github.com/schroeme/AstrocyteCaAnalysis.

## TECHNICAL TERMS

**Astrocytes** are star-shaped glial (non-neuronal) cells of the central nervous system responsible for a variety of functions.

**Gliotransmission** refers to the calcium-dependent release of neurotransmitters, (including glutamate, D-serine, and ATP) from astrocytes that has been demonstrated in some preparations.

**Calcium imaging** is a microscopy technique for measuring optical calcium signals in living cells using fluorescent calcium indicators.

**Genetically-encoded calcium indicators** are used to measure intracellular calcium levels of genetically-defined sub-populations, as they emit fluorescence upon binding to calcium.

**Primary cell culture** refers to cell cultures derived directly from dissociated tissue, as opposed to secondary cultures sub-cultured from other cell cultures.

**Traumatic brain injury** refers to physical, mechanical trauma to neural cells resulting from a blow or jolt to the head or body.

**Glutamate** is a critical neurotransmitter at excitatory synapses and the most abundant free amino acid in the brain.

**Multilayer networks** are a more general network framework which allow for the interaction of multiple node types across layers of a single node type.

