## Supplemental Information for "A multilayer network model of neuron-astrocyte populations in vitro reveals mGluR_5_ inhibition is protective following traumatic injury"

and David F. Meaney[1]

### SUPPORTING INFORMATION

#### *A. Automated detection of neuron and astrocyte calcium events*

Astrocyte calcium events are extremely complex, and there is no stereotypical astrocyte calcium transient, as there is for a neuronal calcium transient or action potential (Bazargani and Attwell (2016); Khakh and McCarthy (2015); Wang et al. (2018, 2017)). The spatial and temporal heterogeneity of astrocyte calcium signals impedes their identification and quantification. Nevertheless, to fully understand neuron-astrocyte signaling, astrocyte $Ca^{2+}$ must be directly observed. We aimed to develop a robust quantitative algorithm for differentiating neuron and astrocyte calcium signals and counting astrocyte calcium events. As with neurons, while some information is lost when a calcium trace is converted to an event train, binarization reduces the complexity of the data and the subsequent computational load. Furthermore, event binarization enables calculation of metrics such as event rate, synchronization, and functional connectivity.

There are several existing methods for the automatic identification of neural spikes from various acquisition modalities (Nenadic and Burdick (2005); Quiroga, Nadasdy, and Ben-Shaul (2004); Schultz, Kitamura, Post-Uiterweer, Krupic, and Häusser (2009)). Our group recently developed FluoroSNNAP, Fluorescence Single Neuron and Network Analysis Package, an open-source, interactive software developed in MATLAB for automatic quantification of single-cell and population-level calcium dynamics (Patel, Man, Firestein, and Meaney (2015)). FluoroSNNAP showed improved calcium transient detection with a template-based algorithm compared to peak-based or wavelet-based detection

Corresponding author: David F. Meaney, methods. Subsequently, we chose to develop an analogous template-based algorithm for automated detection of astrocyte calcium transients. This automated event detection algorithm is integrated into a semi-automated pipeline for analyzing large amounts of calcium image data from mixed cell populations. Here we describe the development of this pipeline and demonstrate its utility in analyzing real astrocyte calcium data.

To generate the astrocyte analog of this neuronal spike detection algorithm, a library of representative astrocyte calcium event templates was curated from actual calcium activity of IF-confirmed astrocytes. Whole traces ($n$ = 60 traces, 180s each) considered to be representative of astrocyte activity were manually selected from the baseline imaging period of all imaged astrocyte segments ($n$ = 5 dishes, $n$ = 125 segments). Traces were smoothed four times with a moving average filter using MATLAB's smooth function. Astrocyte calcium events were detected as local maxima in the scaled flourescence of astrocyte ROIs. The full recording of each smoothed astrocyte calcium fluorescence trace was binned into 60s time windows. Local maxima in each time window were detected using MATLAB's *findpeaks* function, requiring a minimum peak height of 95% of the maximum in that time window, and a minimum distance between peaks of 2s. To prevent false detection of high-frequency, low-amplitude noise, the maximum peak height of each time window was required to be at least as high as the maximum peak of the full trace minus 0.2 in scaled fluorescence. Detected peaks were overlaid on their fluorescence traces and manually sorted. Only peaks that were correctly identified were retained for further analysis. Traces were then broken into snippets of fluorescence activity 1s preceding and following each peak, each snippet being 2s in length and containing one or a few peaks. Snippets were then manually shortened to capture a single peak from its beginning to 75% of its duration (Fig. 12A).

To generalize the waveform library to transients of various widths and heights, library snippets were scaled vertically and horizontally. Widths were scaled to a maximum of 4s and a minimum of 250ms, a reasonable range for the duration of astrocyte calcium transients. Heights were scaled to a maximum of the largest peak of all library traces, and a minimum of the smallest peak of all library traces. 18 horizontal and 100 vertical scaling factors were generated by sampling evenly over the interval from minimum to maximum and normalizing to the mean. Traces were scaled vertically by multiplying the scaled fluorescence by the scaling factor. Horizontal scaling was accomplished by interpolating library traces over every range of horizontal scaling factors using MATLAB's *interp1* function with "pchip," a shape-preserving piecewise cubic interpolation of the values at neighboring grid points. A total of 1,800 scaled versions were generated for 217 library waveforms.

The library of scaled astrocyte calcium waveforms was further pruned to minimize the error between library waveforms and unfiltered astrocyte calcium traces. First, peaks in the original 60 unfiltered scaled fluorescence traces were identified using the same algorithm as for filtered traces. 2s-snippets of unfiltered fluorescence data were created as described above. Mean squared error (MSE) between 280 unfiltered snippets and 217 library snippets (each with 1,800 scaled versions) was calculated as a function of library trace identity and horizontal and vertical scaling factors, using MATLAB's *immse* function. For each of the 280 raw traces, the library trace and corresponding scaling factors that minimized MSE between the two was identified. After removing library waveforms that produced minimum MSE for only one unfiltered trace, 51 best library waveforms remained. MSE calculations were repeated using only these waveforms. To identify the best, as determined by lowest MSE, of these 51 repeated library waveforms, we set a requirement that a library waveform be represented as the best trace for at least two unfiltered traces. The number of variations in vertical and horizontal scaling factors for each waveform was set to equal the number of times it resulted in the minimum MSE for a trace. For example, if if a library waveform produced the best fit for three traces, it was scaled three times and represented three times in the library. This method produced a total of 280 scaled library waveforms generated from 51 parent waveforms, each scaled two or more times.

This library of curated astrocyte waveforms was used as templates in a previously developed template-matching algorithm for neuronal event detection (Patel et al. (2015)). Algorithm parameters were adjusted for analysis of astrocyte, rather than neuron, calcium dynamics. Briefly, this algorithm works as follows. Background fluorescence for all time points is estimated by interpolating from a linear fit to the average background fluorescence, to account for fluctuations in background. Background noise is calculated as five times the standard deviation of the output from a high-pass (order 15, 200 Hz cutoff) Butterworth filter applied to background fluorescence, divided by the mean of the background fluorescence. The raw fluorescence trace is scaled by background fluorescence for subsequent processing. To calculate the signal-to-noise ratio (SNR), noise is defined as one standard deviation above or below the mean scaled fluorescence. To estimate noise, the instantaneous standard deviation of the signal is calculated for 11 different time windows. The SNR in each time window is calculated as the

99th percentile of the instantaneous signal standard deviation divided by the 1st percentile of the instantaneous standard deviation. Overall SNR is defined as the standard deviation of the SNR in each time window (Eq. 1).

$$SNR = \sqrt{\frac{1}{N-1}\sum_{i=1}^{N}\left(\frac{\sigma_{.99,i}}{\sigma_{0.01,i}} - \overline{\frac{\sigma_{.99,i}}{\sigma_{0.01,i}}}\right)^2}, \quad (1)$$

where $N$ is the number of time windows, $\sigma_{0.01,i}$ is the 1st percentile of the instantaneous standard deviation of the $i$th time window, and $\sigma_{0.99,i}$ is the 99th percentile of the instantaneous standard deviation of the $i$th time window. If the SNR is below 1.5, the signal is deemed too low for event detection. Otherwise, the SNR is used to set the threshold for minimum peak level in subsequent peak detection.

To proceed with event detection, each library snippet is slid along the length (in time) of the calcium signal and the Pearson's linear correlation coefficient is calculated at each time point. To eliminate the potential of high correlation between the library waveform and noise, only the correlation of time points where the signal exceeded background noise is recorded. The correlation matrix is then passed through a median filter with a neighborhood size of five. The instantaneous correlation for every library waveform is collapsed into an overall probability of an event (henceforth spike, for illustrative purposes) occurring over all waveforms. Only filtered correlation values above a certain threshold, or below the negative of the threshold, are counted in the high probability and low probability calculations. The high and low spike probability signals are filtered twice more using a 1D median filter with three neighbors and a zero-phase digital filter, and averaged over all library waveforms.

Local maxima in the high and low probability signals are detected, and the mean and maximum value of these peaks in probability are used to create a threshold for further peak detection to avoid false detection of noisy peaks. The algorithm then checks that each peak in the high probability of spike signal is followed and preceded by a peak in the low probability of spike signal. The fine-grained spike time determined to be the first maximum of the gradient of the peaks in the high probability signal. The final step in event detection is manual elimination of falsely identified peaks. Scaled fluorescence traces with detected peaks overlaid are manually inspected by a user. To simplify the analysis protocol, if the majority of the automatically detected peaks are incorrect, the ROI is eliminated from further analysis.

To validate our event detection algorithm, we assessed its performance on 10 recordings from five different isolations, each with one dish and two conditions. We recorded the number of correctly identified events, falsely identified events, and true events. The sensitivity and specificity of event detection were calculated for each trace (Eqs. 2 & 3).

$$Sensitivity = \frac{TP}{TP + FN}, \tag{2}$$

and

$$Specificity = \frac{TN}{TN + FP}, \tag{3}$$

where $TP$ is the number of true events correctly identified, $FN$ is the number of true events missed, $TN$ is the number of windows that were correctly identified as not having an event, and $FP$ is the number of windows incorrectly identified as having an event.

To test the performance of the astrocyte event detection algorithm on fluorescence traces that were not used to generate the library, we assessed its performance on 10 recordings from five different isolations, each with one dish and two conditions. The signal-to-noise ratio for all astrocyte ROIs in four of these recordings fell below the threshold for event detection. A total of 78 astrocyte traces in the remaining six recordings from four dishes were manually inspected. We recorded the number of correctly identified events, falsely identified events, and true events for each trace. We used recording identification as a pooling variable when averaging sensitivity and specificity to avoid bias towards recordings with more traces (more active astrocyte segments). Mean sensitivity of all recordings was 92.61% (95% CI: [0.90, 1.03]) and mean specificity was 96.35% (95% CI: [0.85, 1.00]).

#### *B. A functional assay to differentiate neurons and astrocytes*

Neurons and astrocytes respond differently to the application of N-Methyl-D-aspartic acid or N-Methyl-D-aspartate (NMDA). NMDA, an amino acid derivative, is a specific NMDA receptor (NMDAR) agonist that mimics the action of glutamate, the endogenous NMDAR ligand. Unlike glutamate, NMDA is specific to NMDARs and does not activate other glutamate receptors that may be present on neurons and astrocytes. Evidence of functional NMDAR expression in cultured cortical astrocytes is insufficient to confirm existence (Dzamba, Honsa, and Anderova (2013)). Regardless of expression level, as observed here and in prior studies, NMDA does not directly excite astrocytes,(Backus, Kettenmann, and Schachner (1989); Bowman and Kimelberg (1984); Kettenmann and Schachner (1985); Nagai, Tsugane, Oka, and Kimura (2004)) but has an excitotoxic effect on neurons, greatly increasing their activity. With this knowledge, we can classify cells that exhibit increased calcium event frequency after NMDA application as neurons, and those that are inactive or maintain their basal activity level as astrocytes. Below we describe an experiment we conducted to validate the use of NMDA as a functional terminal assay to distinguish between neurons and astrocytes in our cell culture model.

Primary cultures ($n$=5) prepared as described in the Materials and Methods section were transduced with GCaMP6f on the CAG promoter at DIV 3 and imaged at 488nm, 50s exposure on DIV 7. Three minutes of baseline activity was recorded after a two-minute adjustment period on the stage. Cells were imaged immediately following addition of 100uM NMDA + 1uM glycine coagonist for up to five minutes. Calcium activity was extracted as described in the Materials and Methods (Fig. 9). Maximum fluorescence projections generated in ImageJ software (National Institutes of Health) were manually identified, with neuronal cell bodies and astrocyte microdomains segmented as separate regions of interest (ROIs). Astrocyte segments were labeled using a custom MATLAB graphical user interface). The calcium activity of predicted astrocytes after NMDA application was examined and compared to that of a neuron (Fig. 12C-E). Predicted astrocyte ROIs that responded as a neuron would to NMDA application were reassigned as neurons.

Cells were fixed in 4% PFA and stained per the protocol described below for microtubule-associated protein 2 (MAP2, neurons; 1:1000 dilution for mouse-anti-MAP2 primary and donkey-anti-mouse secondary antibody) and GFAP (astrocytes; 1:500 dilution for rabbit-anti-GFAP primary antibody and 1:1000 dilution for goat-anti-rabbit secondary antibody). DNA was stained with Hoechst at 10ug/mL. Immunoflourescent (IF) images were obtained at 405 (DNA), 561 (neurons), and 640 (astrocytes) nm. To verify that ROIs morphologically and functionally identified as astrocytes expressed GFAP, the same field of view as imaged under GCaMP6f was relocated during IF imaging.

Following the application of NMDA, neuronal event rate increased significantly by a mean of 16.89 events per minute (Fig. 12E, Sidak's multiple comparisons test; $t$ = 4.932, $df$ = 8, $p$ = 0.0003). The effect of NMDA was easily identified by visual examination of single-cell and population activity (Fig. 12D). Importantly, the calcium activity of astrocytes was unaltered from baseline following addition of NMDA (mean difference = -1.493, Sidak's multiple comparisons test; $t$ = 0.5897, $df$ = 8, $p$ = 0.8165). After sorting cells based on NMDA response, 100% of predicted astrocyte segments ($n$ = 79) and 96% of neurons ($n$ = 604) were confirmed by IF staining ($n$ = 5 dishes).

#### *C. Calculation of pairwise correlation for adjacency matrices*

First, the background fluorescence, estimated from the mean fluorescence of 50 non-ROI regions in the field of view, was subtracted from each ROI's fluorescence. The change in fluorescence was scaled to background, and scaled change in fluorescence was filtered using an order 5, 0.5Hz lowpass Butterworth filter, implemented in MATLAB. For a pair of ROIs $x$ and $y$ with time series $x(t)$ and $y(t)$, the Pearson's correlation coefficient (Eq. 4) was calculated over a set of time lags from -1s to 1s:

$$\rho_{xy} = \frac{N \sum x(t)y(t) - (\sum x(t) \sum y(t))}{\sqrt{[N \sum x(t)^2 - (\sum x(t))^2][N \sum y(t)^2 - (\sum y(t))^2]}}, \tag{4}$$

where $N$ is the total number of time points and each sum is taken over $N$.

The maximum value of $\rho_{xy}$ over all time lags was taken and compared to $\rho_{xy}$ between $x(t)$ and 100 surrogate traces $y(t)$, generated using the same AAFT algorithm described above, at the same lag as the actual traces. We then calculated the $Z$-statistic for the maximum actual $\rho_{xy}$ based on the distribution of $\rho_{xy}$'s generated using the surrogates, and converted it to a $p$-value based on the standard normal distribution. The $p$-value is the probability that the observed $\rho_{xy}$ came from a distribution of $\rho_{xy}$'s between $x(t)$ and a randomly permuted $y(t)$ with the same frequency and amplitude spectra. If $p$ was less than 0.001, $A_{xy}$, the weight of the edge between astrocyte segments $x$ and $y$, was set equal to the maximum value of $\rho_{xy}$, and zero otherwise.

#### *D. Network Statistics*

Mean degree is the mean number of edges emerging from each node, or number of other nodes to which each node is connected. It is defined mathematically as:

$$\langle K \rangle = \frac{1}{N} \sum_{i=1}^{N} A_{ij}, \tag{5}$$

where $A_{ij}$ is the binary weight between nodes $i$ and $j$, and where $N$ is the total number of nodes in the network. A high value of $\langle K \rangle$, which we normalize to $N$, means that on average, nodes in the network are connected to a large proportion of other nodes in the network.

Density is the ratio of the number of actual edges in the network to the total number of potential edges in the network. The number of potential edges in the network is proportional to the number of nodes: each node can be connected to each of the other nodes, but not itself. It is defined mathematically as:

$$\kappa = \frac{R}{N(N-1)}, \tag{6}$$

where $R$ is the total number of edges in the network. Conceptually, density is an indicator of how strongly connected the network is. A network with high density has strongly interconnected nodes, while a network with low density has less strongly interconnected nodes.

Nodal strength is similar to degree, but accounts for connection weight in weighted graphs. Nodal strength is the sum of the weights of all of a node's edges in a weighted network. Conceptually, strength can be thought of as a weighted degree. It is defined mathematically as:

$$S(i) = \frac{1}{N-1} \sum_{j=1}^{N} A_{ij}. \tag{7}$$

In this work, we calculated the average network strength over all nodes, normalized by network size $N$. It is important to normalize by $N$ because larger networks have a greater number of weighted connections and therefore a higher upper bound for $\langle S \rangle$. A network with high normalized $\langle S \rangle$ has many nodes with either many connections (high degree), a number of strong connections (large weight values), or both, for its size. Conversely, a network with low normalized $\langle S \rangle$ has many nodes with either few connections (low degree), a number of weak connections (low weight values), or both, for its size.

Mean clustering coefficient is the mean ratio of the number of actual triangles around each node to the total number of potential triangles around each node. It measures how many sets of three nodes are fully interconnected, or where each of the three nodes is connected to the other two, and thus can be conceptualized as a local density measure. It is defined mathematically as:

$$C = \frac{1}{n} \sum_{i=1}^{N} \frac{2u_i}{k_i(k_i-1)}, \tag{8}$$

where $k_i$ is the degree of node $i$ and $u_i$ is the number of triangles containing node $i$. The clustering coefficient is generalized to weighted networks by replacing the number of triangles $u_i$ with the sum of triangle intensities (Onnela, Saramäki, Kertész, and Kaski (2005)). The averaged weighted clustering coefficient is calculated as:

$$C = \frac{1}{n}\sum_{i=1}^{N}\frac{2}{k_i(k_i-1)}\sum_{j,k}(\tilde{w_{ij}}\tilde{w_{jk}}\tilde{w_{ki}})^{1/3}, \tag{9}$$

where the weights are scaled by the largest weight in the network, $\tilde{w_{ij}} = \tilde{w_{ij}}/max(w_{ij})$. Conceptually, by this definition, a node's weighted clustering coefficient is the unweighted version renormalized by the average intensity of triangles at that node. Here, we calculated the mean clustering coefficient, $\langle C \rangle$, in each of our networks. A network with a high value of $\langle C \rangle$ has a large proportion of fully connected triangles, or high local density, while a network with a low value of $\langle C \rangle$ has a small proportion of fully connected triangles, or low local density.

Betweenness centrality is a measure of how many shortest path lengths pass through a given node. Conceptually, betweenness centrality measures the degree to which a node acts as a hub, facilitating many shortest-path connections between other nodes. It is defined as:

$$B_i = \frac{1}{(n-1)(n-2)}\sum_{h\neq j, h\neq i, j\neq i}^{N}\frac{l_{hj}(i)}{l_{hj}}, \tag{10}$$

where $\rho_{hj}$ is the number of shortest paths between nodes $h$ and $j$, and $l_{hj}(i)$ is the number of shorstest paths between node $h$ and node $j$ that pass through node $i$. The shortest path length between nodes $i$ and $j$ is the minimum number of nodes that must be passed through to connect nodes $i$ and $j$. For weighted networks, betweenness centrality was calculated as above, using the weighted distance matrix, computed using the BCT's distance_wei function, which uses Dijkstra's shortest path algorithm. Brandes's algorithm (Brandes (2001)) was used to compute centrality from the weighted distance matrix via the BCT's *betweenness_wei* function. We calculated the mean nodal betweenness centrality, $\langle B \rangle$, for each of our networks. We normalized by the number of nodes in the network $N$ because a larger network has a greater number of shortest paths and therefore a larger upper bound on $\langle B \rangle$. A network with a high value of normalized $\langle B \rangle$ has a large proportion of nodes that act as hubs, with several shortest paths passing through them, while a network with a small value of $\langle B \rangle$ has a low proportion of hub-like nodes.

Global efficiency is the inverse of the harmonic mean of the shortest path length between any two nodes. The name refers to the fact that a network with a high characteristic shortest path length will, under some specific assumptions of dynamics, be slower to transmit information from node $i$ to node $j$ than a network with many short paths between nodes. Networks with few long-distance connections typically have many large shortest path lengths. Reaching node $j$ from node $i$ in such a network requires many steps through other nodes, lessening its supposed efficiency. Global efficiency is defined as:

$$E = \frac{1}{n} \sum_{i=1}^{n} \frac{\sum_{i \neq j}^{N} d_{ij}^{-1}}{n-1}, \tag{11}$$

where $d_{ij}$ is the shortest topological distance between nodes $i$ and $j$. Global efficiency of weighted networks is calculated as above with $d_{ij}$ being a weighted path length, calculated using Dijkstra's algorithm in the BCT's efficiency_wei. As described above, a network with a high value of $E$ has many long topological connections, a short characteristic pathlength, and is faster, under some specific assumptions of the dynamics, to transmit information between any two nodes. Conversely, a network with a low value of $E$ has few long topological connections, a large characteristic pathlength, and is slower, under some specific assumptions of the dynamics, to transmit information between any two nodes.

#### *E. Contribution of relative abundance of neurons to multilayer network topology and community structure*

To determine the impact on micro- and macroscale multilayer topology of there being many more active neurons than astrocyte segments, we sub-sampled the neuronal population so that neuron and astrocyte layers were of equal size. For a multilayer network with $n_a$ active astrocyte segments, we randomly selected $n_a$ neurons from the neuronal population and formed new adjacency matrices based on the connectivity of the sub-population of neurons and the original population of astrocyte segments. Mean nodal strength, degree, density, betweenness centrality, clustering coefficient, and global efficiency were re-calculated for the half-neuron, half-astrocyte networks. Likewise, we re-ran community detection analysis and re-calculated the $ARI$ and cell type module participation for balanced networks. Neuron layers were sub-sampled 30 times and the average measures over 30 iterations are reported. While this analysis is not biologically realistic, it is a useful statistical excercise to determine the impact of the relative abundance of active neurons in our cultures.

As was found for full-sized multilayer networks, strength in neuron and astrocyte layers (one-sample $t$-test on differences between actual values and null values of $S$: N-N, $t$ = 2.258, $df$ = 21, $p$ = 0.0347; A-A, $t$ = 4.554, $df$ = 21, $p$ = 0.0002) was higher than randomized controls, while mean nodal strength between layers was lower than randomized controls (Fig. S6A, one-sample $t$-test on differences between actual values and null values of $S$, $t$ = 3.795, $df$ = 21, $p$ = 0.0011). Thus, as with full-sized multilayer networks, intralayer connections were stronger than interlayer connections. This finding suggests that the relative strength of intralayer connections compared to interlayer connections is not primarily driven by the greater abundance of neurons. However, astrocyte segments were no longer significantly more strongly interconnected than neurons (Tukey's multiple comparisons test following one-way ANOVA, $q$ = 2.731, $df$ = 63, $p$ = 0.1384), suggesting that differences in mean normalized nodal strength between layers are primarily due to layer size, to which strength is normalized. As was found for actual multilayer networks, randomly sub-sampled multilayer networks with an equal number of neurons and astrocytes exhibited significantly larger clustering coefficient and lower global efficiency than their randomized counterparts (Fig. 6B), but were not different in betweenness centrality $B$. Topological measures $C$, $B$, and $E$ were similarly correlated with density in sub-sampled multilayer networks (Fig. S6C).

As was done for full-sized multilayer networks, we analyzed the impact of experimental manipulations on sub-sampled multilayer network topology using a generalized linear regression, with mean nodal strength as a regressor (Table S 7). In sub-sampled multilayer networks, no manipulation was a significant predictor of topology after controlling for multilayer strength. This finding suggests that injury-mediated changes in clustering coefficient are primarily driven by the relative abundance of neurons in our cultures. Furthermore, in sub-sampled multilayer networks, neither treatment with MPEP, injury, or their interaction was a significant predictor of multilayer or interlayer mean nodal strength at the final experimental time point (GLM with $S$ as outcome variable and MPEP, Injury, MPEP + Injury interaction term, and event rate as covariates, all $p$-values from $z$-tests on coefficients above 0.05).

To assess the impact of the relative abundance of neurons in our cultures on community structure, we performed the same modularity detection on sub-sampled multilayer networks ($\gamma_s = 1.13, \gamma_f = 1.55$). The mean $ARI$ between spatial and functional sub-sampled multilayer communities was significantly larger than for actual multilayer networks (mean for full multilayer = 0.05364, mean for sub-sampled multilayer = 0.1065, paired two-tailed $t$-test, $t$ = 2.346, $df$ = 22, $p$ = 0.0284). This disagreement between spatial and functional partitioning was not driven by differences in community size, as $\gamma$, the spatial tuning parameter, was adjusted to reduce this difference (Fig. S6D, paired $t$-test, $t$ = 0.8896, $df$ = 22, $p$ = 0.3833). The difference between actual and sub-sampled multilayer $ARI$ may be due to differences in overall network size, or higher disagreement between spatial and functional modularity in neuronal layers. In sub-sampled multilayer networks, neurons and astrocyte segments participated equally in spatial and functional modules (Fig. S6E). Thus, differences in module participation in full multilayer networks reflects the dominance of neurons in quantity.

#### *F. Community detection methodology*

The modularity quality function is given by:

$$Q = \sum_{ij} [A_{ij} - \gamma P_{ij}] \delta(c_i, c_j), \tag{12}$$

where $Q$ measures quality, $A_{ij}$ is the observed adjacency matrix, $P_{ij}$ is the null model adjacency matrix, and $\delta(c_i, c_j)$ is 1 when nodes $i$ and $j$ are in the same community and 0 otherwise. The parameter $\gamma$ is a resolution parameter that governs the size and number of detected communities. The values of $\gamma$ were 1.125 for spatial astrocyte graphs, 1.270 for functional astrocyte graphs, 1.05 for spatial neuron graphs, 1 for functional neuronal graphs, 1.13 for spatial multilayer graphs, and 1.55 for functional multilayer graphs. A Newman-Girvan (Newman and Girvan (2004)) null model was used for both functional and spatial graphs and it was implemented in a custom MATLAB script using Eqs. 13 - 15:

$$P_{ij} = \frac{s_i s_j}{2m}, \tag{13}$$

where

$$s_i = \sum_j A_{ij}, \tag{14}$$

and

$$m = \frac{1}{2} \sum_{ij} A_{ij}, \tag{15}$$

where $s_i$ is the strength of node $i$, the sum of all its weights to other nodes.

The modularity quality function was maximized using a Louvain-like community detection algorithm. Briefly, the Louvain algorithm is a greedy optimization algorithm wherein each nodes starts in its own community, node community assignment is changed locally, and changes that increase modularity are kept until there are no further increases in quality (Blondel, Guillaume, Lambiotte, and Lefebvre (2008); Fortunato (2010); Mucha, Richardson, Macon, Porter, and Onnela (2010); Porter, Onnela, and Mucha (2009)). The Louvain algorithm was iterated until the algorithm converged on the final module assignment. Because the modularity landscape is rough, with many near-optimal solutions (Good,

De Montjoye, and Clauset (2010)), we assigned a node's module to be the mode of its module assignment over 50 optimizations.

For both single-layer and multilayer networks, $\gamma$ was tuned separately and manually for spatially and functionally generated graphs to minimize the difference between the number of communities detected for spatial and functional networks. In addition to the value of $Q$, the Louvain algorithm outputs the partition $g$, a vector containing the community number of each node. The Adjusted Rand Index ($ARI$) was used to measure the similarity of community partitions. It is defined mathematically as:

$$ARI = \frac{\binom{N}{2}(a+f) - [(a+b)(a+e)+(e+f)(b+f)]}{\binom{N}{2}^2 - [(a+b)(a+e)+(e+f)(b+f)]}, \tag{16}$$

where $N$ is the total number of nodes, $a$ is the number of pairs of nodes that are in the same community in the functional partition, $g_f$, and the spatial partition, $g_s$, $b$ is the number of pairs of nodes that are in a different community in $g_f$ and $g_s$, $e$ is the number of pairs of nodes that are in the same subset in $g_f$ and a different subset in $g_s$, and $f$ is the number of pairs of nodes that are in a different subset in $g_f$ and the same subset in $g_s$. For astrocytes, $ARI$ was also calculated for $g_a$, the actual partitioning of astrocyte segments into cells, versus $g_f$ and $g_s$. The Adjusted Rand Index can be negative, indicating that the partitions disagree more than what would be predicted by chance, and has a maximum of 1 for total agreement.

#### *G. Immunocytochemistry*

Cells were fixed in 4% paraformaldehyde (PFA) immediately following imaging and maintained in 1X phosphate-buffered saline (PBS) until staining. Cell membranes were permeabilized with cold 0.2% Triton X-100 in PBS for 5 min. Non-specific binding was blocked with 1% bovine serum albumin (BSA, Sigma) and 2.5% normal goat serum (NGS) for 45 min at room temperature (RT). Primary antibodies were incubated overnight at 4°C in diluted blocking solution (0.2% BSA and 0.5% NGS in PBS) at the following concentrations: Mouse-anti-MAP2 (Millipore Sigma) at 1:750, Rabbit-anti-GFAP (Abcam) at 1:500. Following a wash with diluted blocking solution, secondary antibodies were incubated in diluted blocking solution for 45 min at RT at the following concentrations: Goat-anti-mouse Alexa Fluor 568 (Thermo Fisher Scientific) at 1:1000 and Goat-anti-rabbit Alexa Fluor 633 (Thermo Fisher Scientific) at 1:1000. After a second wash, anti-mGluR5 Alexa Fluor 488 (Novus Biologicals) was incubated at a concentration of 1:100. All antibody solutions were centrifuged at 15,000 rotations per minute (RPM) for 10 min to remove aggregates. Following three rinses with PBS, stained cells were maintained under low light conditions until imaging. To stain for nuclei, 10 ug/mL Hoescht was applied during the final rinse step.

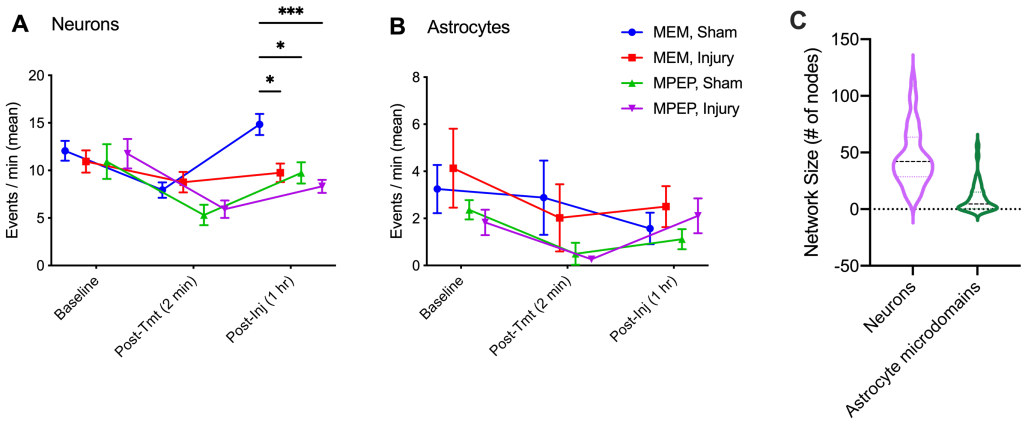


**Supplementary Figure 1.** Calcium activity and size of neuron-astrocyte cultures. **A-B.** $mGluR_5$ inhibition and injury decrease neuronal but not astrocytic activity level. **A.** Neuronal event rate (events/min) at the three measured time points for all four treatment groups (n = 9 dishes for MEM/Sham and MEM/Inj, n = 8 for MPEP/Sham, n = 10 for MPEP/Inj). **B.** Astrocytic event rate (events/min) at the three measured time points. Error bars indicate standard error of the mean (SEM) and asterisks indicate statistical significance (*$p \leq 0.05$, **$p \leq 0.01$, ***$p \leq 0.001$, ****$p \leq 0.0001$). Tmt: treatment; MEM: treated with minimum essential media; MPEP: treated with anti-$mGluR_5$; Injury: subjected to targeted neuronal tap injury; Sham: negative injury control. **C.** Number of active neurons (purple) or astrocyte microdomains (green) in each dish (95% of number of neurons [37.16, 54.34], 95% CI of number of astrocyte microdomains [5.516, 14.43]). Violin plots show frequency distribution of the data, with dotted lines indicating median and quartiles.

| | Neuronal | | | Astrocytic | | |
|---|---|---|---|---|---|---|
| | diff, 95% CI | $q$ | $p$ | diff, 95% CI | $q$ | $p$ |
| MEM, Sham vs. MEM, Injury | [0.8790, 9.293]* | 4.470 | 0.0111 | [-4.518, 2.661] | 0.9619 | 0.9043 |
| MEM, Sham vs. MPEP, Sham | [0.7588, 9.432]* | 4.345 | 0.0145 | [-3.269, 4.180] | 0.4549 | 0.9884 |
| MEM, Sham vs. MPEP, Injury | [2.409, 10.61]* | 5.869 | 0.0004 | [-3.940, 2.860] | 0.5908 | 0.9753 |
| MEM, Injury vs. MPEP, Sham | [-4.327, 4.346] | 0.007942 | $>$0.9999 | [-2.205, 4.973] | 1.434 | 0.7418 |
| MEM, Injury vs. MPEP, Injury | [-2.678, 5.524] | 1.283 | 0.8010 | [-2.863, 3.639] | 0.4441 | 0.9892 |
| MPEP, Sham vs. MPEP, Injury | [-2.820, 5.647] | 1.235 | 0.8187 | [-4.396, 2.405] | 1.089 | 0.8677 |

**Supplementary Table 1.** Results of Tukey's multiple comparisons test following a 2-way ANOVA on the effect of time and group assignment on neuronal and astrocytic event rate at the final experimental time point (one hour post-injury). For neuron event rate, the results of the 2-way ANOVA were as follows: Time x Group factor, F(6, 64) = 2.738, p = 0.0197; Time factor, F(2, 64) = 22.40, p $<$ 0.0001; Group factor, F(3, 32) = 3.081, p = 0.0412. For astrocyte event rate, the results of the 2-way ANOVA were as follows: Time x Group factor, F(6, 48) = 0.7730, p = 0.5950; Time factor, F(2, 48) = 3.265, p = 0.0468; Group factor, F(3, 24) = 1.604, p = 0.2147. Asterisks indicate statistical significance.

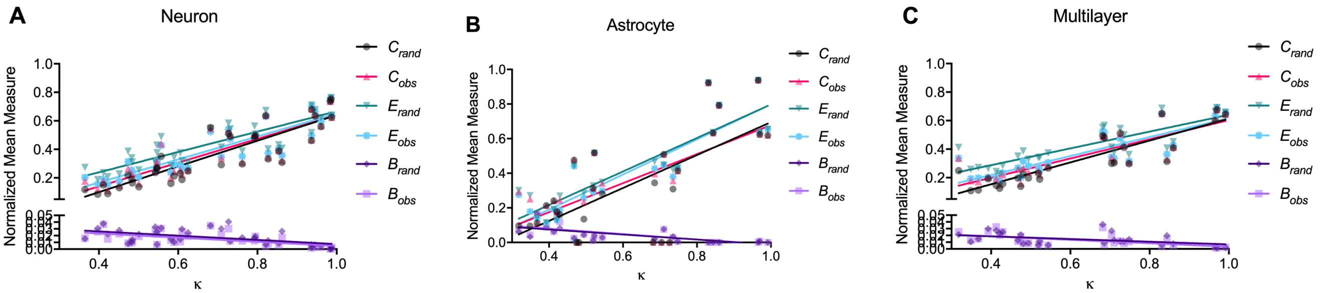


**Supplementary Figure 2.** Dependence of observed and randomized network topology on edge density. **A-C.** $B$, $C$, and $E$ vs network density $\kappa$ for observed (obs) and randomized (rand) networks at the final experimental time point (1 hour post-injury) for neuron-neuron **A**, astrocyte-astrocyte **B** and multilayer networks **C**. See Fig. S2 for statistical details.

**A. Neurons**

| | $B_{rand}$ | $B_{obs}$ | $C_{rand}$ | $C_{obs}$ | $E_{rand}$ | $E_{obs}$ |
|---|---|---|---|---|---|---|
| **95% CI of slope** | -0.04847 to -0.01428 | -0.04529 to -0.01631 | 0.7341 to 1.069 | 0.6659 to 0.9998 | 0.5650 to 0.8608 | 0.6591 to 0.9539 |
| **$R^2$** | 0.2904 | 0.3544 | 0.7791 | 0.7514 | 0.7383 | 0.7843 |
| **F** | 13.92 | 18.67 | 119.9 | 102.7 | 95.91 | 123.6 |
| **DF** | 34 | 34 | 34 | 34 | 34 | 34 |
| **p** | 0.0007 | 0.0001 | <0.0001 | <0.0001 | <0.0001 | <0.0001 |

**B. Astrocytes**

| | $B_{rand}$ | $B_{obs}$ | $C_{rand}$ | $C_{obs}$ | $E_{rand}$ | $E_{obs}$ |
|---|---|---|---|---|---|---|
| **95% CI of slope** | -0.2226 to -0.06700 | -0.2359 to -0.07844 | 0.5367 to 1.366 | 0.4147 to 1.264 | 0.6867 to 1.251 | 0.7334 to 1.296 |
| **$R^2$** | 0.4298 | 0.4644 | 0.4719 | 0.3987 | 0.7085 | 0.7285 |
| **F** | 15.07 | 17.34 | 22.34 | 16.58 | 51.05 | 56.34 |
| **DF** | 20 | 20 | 25 | 25 | 21 | 21 |
| **p** | 0.0009 | 0.0005 | <0.0001 | 0.0004 | <0.0001 | <0.0001 |

**C. Multilayer**

| | $B_{rand}$ | $B_{obs}$ | $C_{rand}$ | $C_{obs}$ | $E_{rand}$ | $E_{obs}$ |
|---|---|---|---|---|---|---|
| **95% CI of slope** | -0.03650 to -0.003395 | -0.04029 to -0.01069 | 0.5965 to 0.9362 | 0.5049 to 0.8436 | 0.4294 to 0.7477 | 0.5048 to 0.8263 |
| **$R^2$** | 0.1977 | 0.3347 | 0.7755 | 0.729 | 0.6989 | 0.7442 |
| **F** | 6.16 | 12.58 | 86.35 | 67.24 | 58.02 | 72.74 |
| **DF** | 25 | 25 | 25 | 25 | 25 | 25 |
| **p** | 0.0201 | 0.0016 | <0.0001 | <0.0001 | <0.0001 | <0.0001 |

**Supplementary Table 2.** Results of linear regressions of $B$, $C$, and $E$ on network density $\kappa$ for observed (obs) and randomized (rand) networks at the final experimental time point (1 hour post-injury, see Table S2). Reported are the 95% confidence interval on the slope, the $R^2$ value, F-statistic, degrees of freedom (DF), and the p-value for neuron-neuron networks (**A**), astrocyte-astrocyte networks (**B**) and multilayer networks (**C**). For all network types, a stronger correlation was seen between $B_{obs}$ and $E_{obs}$ and $kappa$ than for $B_{rand}$ and $E_{rand}$ and $kappa$, suggesting that these aspects of topology are more dependent on edge density for our networks than would be expected at random.

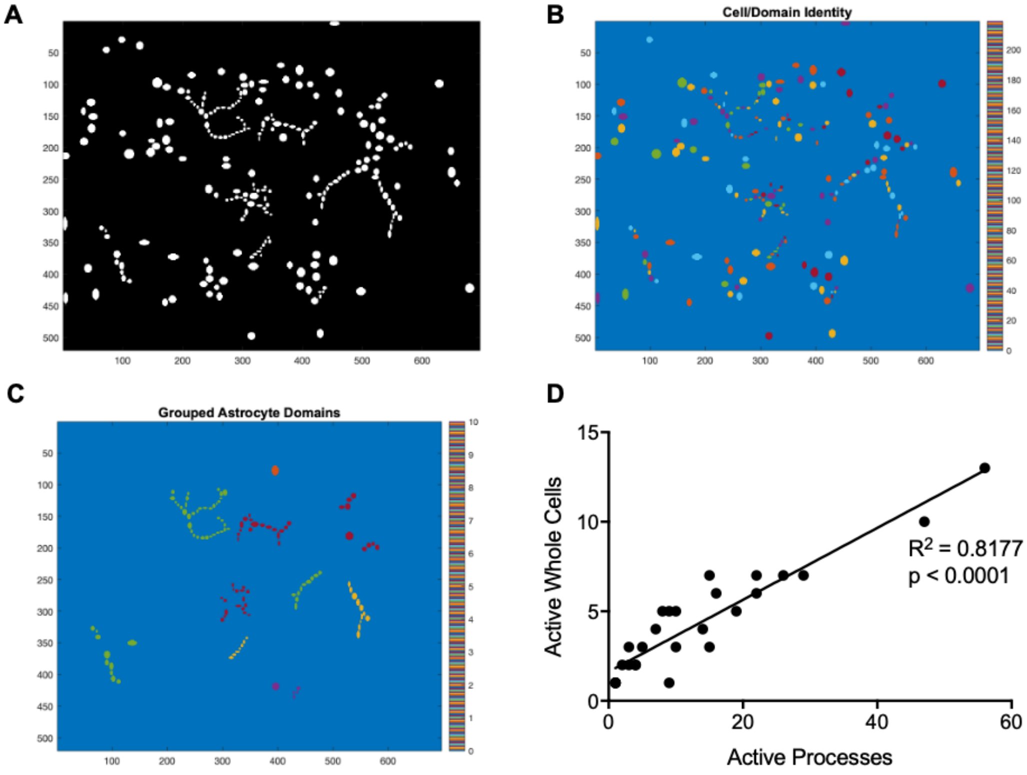


**Supplementary Figure 3.** Manual grouping of astrocyte microdomains. **A.** Binary segmentation mask used to extract continuous calcium signal for a representative field of view. Neurons and astrocyte domains are indicated as nonzero (white) pixels. **B.** Mask colored by cell or astrocyte microdomain identity (index 1-217), before grouping of astrocyte segments. **C.** Mask of astrocyte domains only, grouped by manually-identified cell using a custom-built astrocyte identification graphical user interface (GUI). The GUI allows users to upload a segmentation file and click on predicted astrocyte segments to label them as such for downstream image analysis. Users can remove falsely labeled astrocytes in the GUI. Duplicates are automatically removed. **D.** Number of active whole astrocytes vs. number of active astrocyte microdomains (processes). As expected, there is significant correlation between the two (simple linear regression, 95% CI of slope [0.1681, 0.2336], $R^2 = 0.8643$, $F(1,25) = 159.3$, $p < 0.0001$.

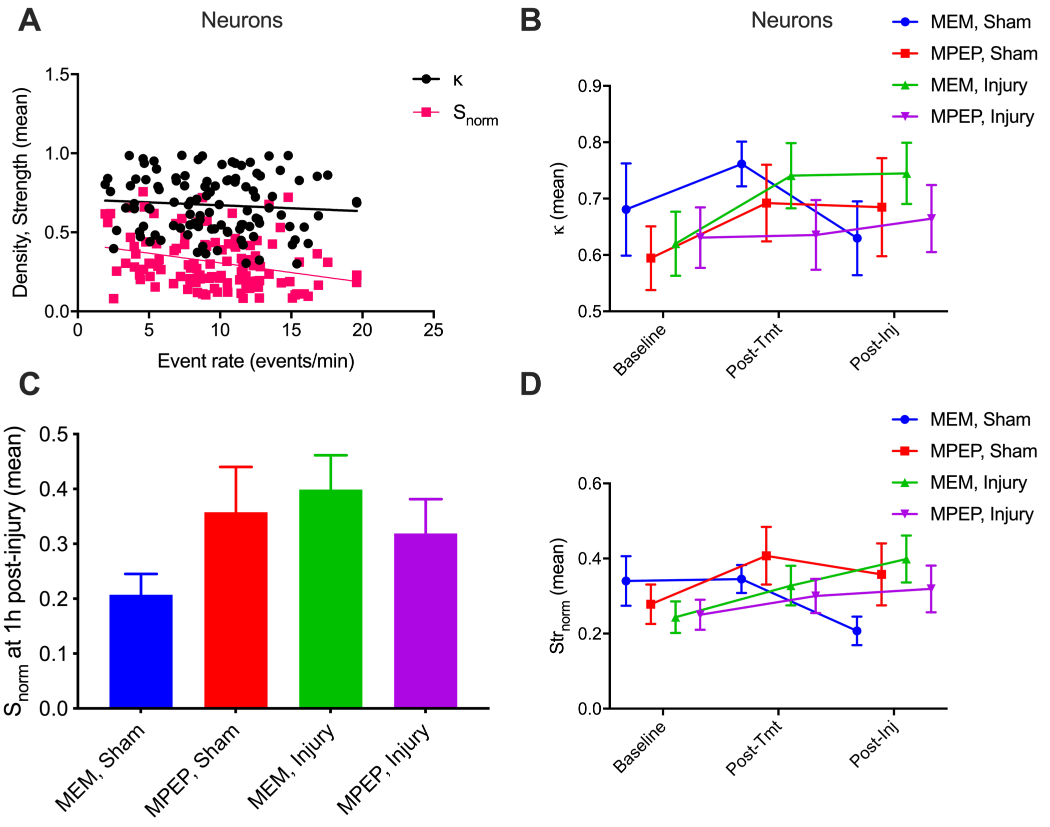


**Supplementary Figure 4.** Effect of exogenous manipulations on neuron network edge density and nodal strength. **A.** Neuron-neuron network mean edge density (black) and mean normalized nodal strength (magenta) versus event rate for all dishes at all experimental time points (simple linear regressions: $\kappa$, 95% CI of slope [-0.01256, 0.005102], $F$(1,106) = 0.7007, $R^2$ = 0.006567, $p$ = 0.4044; $S$, 95% CI of slope [-0.01996, -0.004531], $F$(1,106) = 9.902, $R^2$ = 0.08543, $p$ = 0.0021). **B.** Neuron-neuron network edge density at each experimental time point for each experimental group. **C.** Mean normalized nodal strength for all experimental groups at the final time point, 1 hour post-injury. The differences between groups were not significant (ordinary one-way ANOVA, p = 0.1782). **D.** Mean normalized nodal strength for all experimental groups at all experimental time points. Error bars indicate standard error of the mean (SEM) and asterisks indicate statistical significance (no asterisks, ns, *p $\leq$ 0.05, **p $\leq$ 0.01, ***p $\leq$ 0.001, ****p $\leq$ 0.0001). Tmt: treatment; MEM: treated with minimum essential media; MPEP: treated with anti-mGluR$_5$; Injury: subjected to targeted neuronal tap injury; Sham: negative injury control.

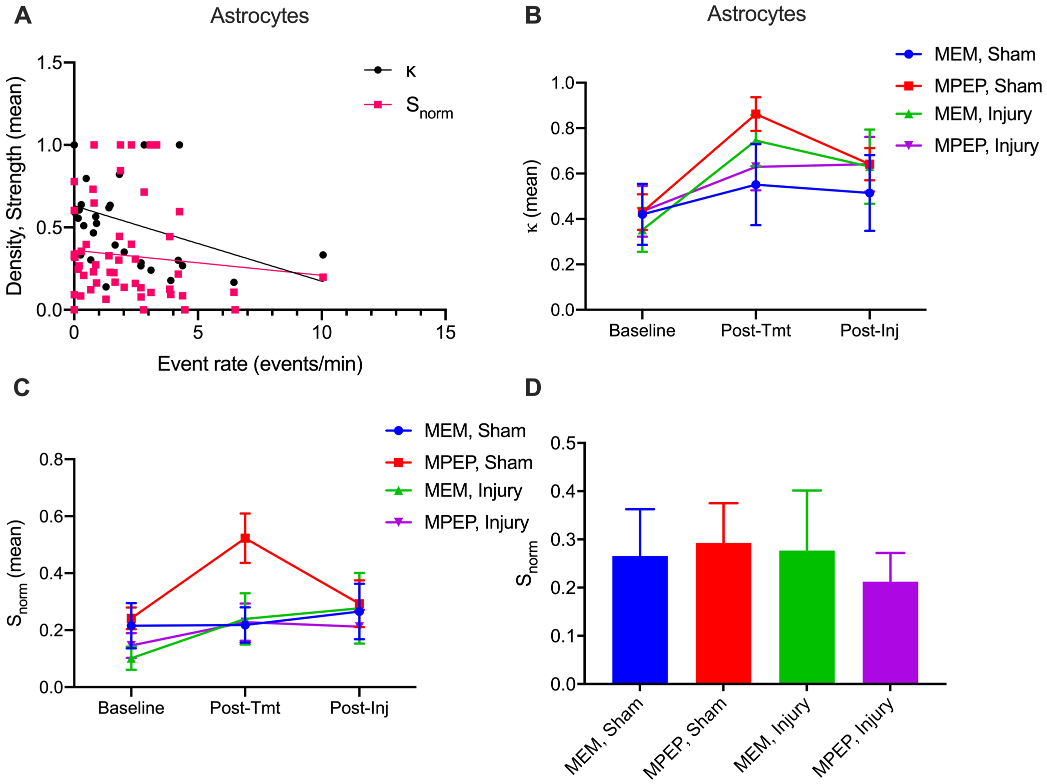


**Supplementary Figure 5.** Effect of exogenous manipulations on astrocyte network edge density and nodal strength. **A.** Astrocyte-astrocyte network mean edge density (black) and mean normalized nodal strength (magenta) versus event rate for all dishes at all experimental time points (simple linear regressions: $\kappa$, 95% CI of slope [-0.07971, -0.01170], $F(1{,}73) = 7.174$, $R^2 = 0.08948$, $p = 0.0091$; $S$, 95% CI of slope [-0.04667, 0.01627], $F(1{,}73) = 0.9267$, $R^2 = 0.01254$, $p = 0.3389$). **B.** Astrocyte-astrocyte network edge density at each experimental time point for each experimental group. **C.** Mean normalized nodal strength for all experimental groups at the final time point, 1 hour post-injury. The differences between groups were not significant (ordinary one-way ANOVA, p = 0.9221). **D.** Mean normalized nodal strength for all experimental groups at all experimental time points. Error bars indicate standard error of the mean (SEM) and asterisks indicate statistical significance (no asterisks, ns, *p $\leq$ 0.05, **p $\leq$ 0.01, ***p $\leq$ 0.001, ****p $\leq$ 0.0001). Tmt: treatment; MEM: treated with minimum essential media; MPEP: treated with anti-mGluR$_5$; Injury: subjected to targeted neuronal tap injury; Sham: negative injury control.

| | $C$ | | | $B$ | | | $E$ | | |
|---|---|---|---|---|---|---|---|---|---|
| | $\beta$ | z | p | $\beta$ | z | p | $\beta$ | z | p |
| Intercept | 0.0815* | 5.783 | 0.000 | 0.0237* | 5.482 | 0.000 | 0.1116* | 11.023 | 0.000 |
| Strength | 0.9306* | 35.137 | 0.000 | -0.0316* | -3.883 | 0.000 | 0.8970* | 47.144 | 0.000 |
| MPEP | 0.0099 | 0.761 | 0.446 | 0.0025 | 0.620 | 0.535 | 0.0042 | 0.452 | 0.652 |
| Sham | -0.0346* | -2.450 | 0.014 | -0.0020 | -0.471 | 0.638 | *-0.0224 | -2.207 | 0.027 |
| MPEP + Sham | 0.0270 | 1.372 | 0.170 | 0.0033 | 0.542 | 0.588 | 0.0174 | 1.234 | 0.217 |

**Supplementary Table 3.** Results of generalized linear regression to predict the effect of group assignment on mean clustering coefficient ($C$), mean normalized betweenness centrality ($B$), and global efficiency ($E$) for **neuron** networks at the final experimental timepoint. $\beta$: estimated coefficient, $z$: value of test statistic for coefficient, the value of the estimate divided by the standard error of the estimate, and $p$: p-value for coefficient resulting from a t-test, pr($> z$), $df$ = 31. The $z$-test tests the null hypothesis that the coefficient for that covariate is equal to zero. Asterisks indicate statistical significance ($p < 0.05$). In this case, only changes in $C$ and $E$ were significantly predicted by injury alone (see effect of Sham).

| | $C$ | | | $B$ | | | $E$ | | |
|---|---|---|---|---|---|---|---|---|---|
| | $\beta$ | z | p | $\beta$ | z | p | $\beta$ | z | p |
| Intercept | *0.1383 | 3.611 | 0.000 | 0.1057* | 4.563 | 0.000 | 0.0844* | 2.128 | 0.033 |
| Strength | 1.0433* | 15.711 | 0.000 | -0.1741* | -4.335 | 0.000 | 1.1275* | 16.392 | 0.000 |
| MPEP | -0.0355 | -0.936 | 0.350 | -0.0237 | -1.033 | 0.302 | -0.0229 | -0.584 | 0.559 |
| Sham | -0.0015 | -0.037 | 0.917 | 0.0082 | 0.336 | 0.737 | 0.0089 | 0.212 | 0.832 |
| MPEP + Sham | -0.0359 | -0.653 | 0.514 | 0.0166 | 0.499 | 0.618 | -0.0334 | -0.587 | 0.557 |

**Supplementary Table 4.** Results of generalized linear regression to predict the effect of group assignment on mean clustering coefficient ($C$), mean normalized betweenness centrality ($B$), and global efficiency ($E$) for **astrocyte** networks at the final experimental time point. $\beta$: estimated coefficient, $z$: value of test statistic for coefficient, the value of the estimate divided by the standard error of the estimate, and $p$: p-value for coefficient resulting from a t-test, pr($> z$), $df$ = 17. The $z$-test tests the null hypothesis that the coefficient for that covariate is equal to zero. In this case, no topological parameters could be significantly predicted by treatment condition.

| | $C$ | | | $B$ | | | $E$ | | |
|---|---|---|---|---|---|---|---|---|---|
| | $\beta$ | z | p | $\beta$ | z | p | $\beta$ | z | p |
| Intercept | 0.1105* | 7.908 | 0.000 | 0.0195* | 4.446 | 0.000 | 0.1252* | 9.461 | 0.000 |
| Strength | 0.8564* | 29.945 | 0.000 | -0.0262* | -2.926 | 0.003 | 0.8457* | 31.211 | 0.000 |
| MPEP | -0.0056 | -0.424 | 0.672 | 0.0032 | 0.767 | 0.443 | -0.0017 | -0.134 | 0.894 |
| Sham | -0.0368* | -2.473 | 0.013 | -0.0008 | -0.162 | 0.871 | -0.0208 | -1.474 | 0.141 |
| MPEP + Sham | 0.0359 | 1.717 | 0.086 | -0.0008 | -0.117 | 0.907 | 0.0196 | 0.991 | 0.321 |

**Supplementary Table 5.** Results of generalized linear regression to predict the effect of group assignment on mean clustering coefficient ($C$), mean normalized betweenness centrality ($B$), and global efficiency ($E$) for **multilayer** networks at the final experimental timepoint. $\beta$: estimated coefficient, $z$: value of test statistic for coefficient, the value of the estimate divided by the standard error of the estimate, and $p$: p-value for coefficient resulting from a t-test, pr($> z$), $df$ = 22. The $z$-test tests the null hypothesis that the coefficient for that covariate is equal to zero. In this case, only changes in $C$ were significantly predicted by injury alone (see effect of Sham).

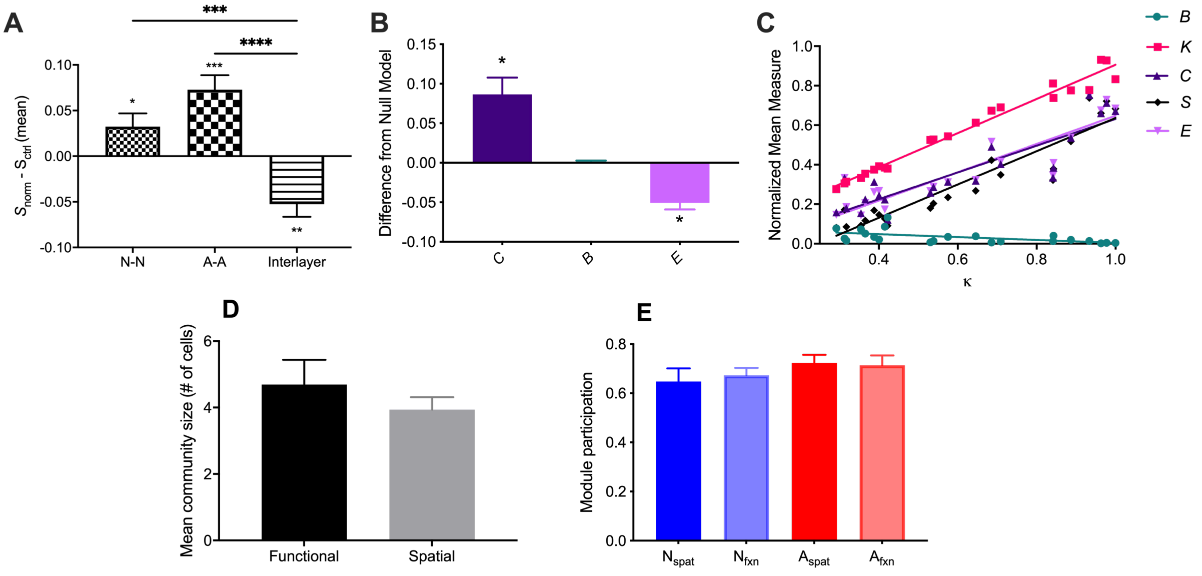


**Supplementary Figure 6.** Characterization of sub-sampled multilayer neuron-astrocyte functional and spatial network topology. Neurons were randomly sub-sampled so that the number of neurons and astrocyte microdomains was equal. **A.** Difference in mean normalized strength of neuron layer, astrocyte layer, interlayer, and multilayer connections of sub-sampled multilayer networks, compared to a randomized network with preserved degree distribution and density (one-sample $t$-tests on difference from randomized networks: N-N, $t$ = 2.258, $df$ = 21, $p$ = 0.0347; A-A, $t$ = 4.554, $df$ = 21, $p$ = 0.0002; N-A, $t$ = 3.795, $df$ = 21, $p$ = 0.0011; Tukey's multiple comparisons test following one-way ANOVA, N-N vs. A-A, $q$ = 2.731, $df$ = 63, $p$ = 0.1384). **B.** Difference from random null model of calculated mean clustering coefficient $C$, normalized betweenness centrality $B$, and global efficiency $E$. We observe significantly larger clustering coefficients (one sample $t$-test, $t$ = 4.103, $df$ = 21, $p$ = 0.0005) and significantly lower global efficiency (one sample $t$-test, $t$ = 6.089, $df$ = 21, $p < 0.0001$) than expected from a random null model with preserved degree distribution. **C.** Mean clustering coefficient $C$, normalized betweenness centrality $B$, normalized degree $K$, normalized strength $S_{norm}$, and global efficiency $E$ vs. mean density $\kappa$ for each dish at the third imaging time point (1 hour post-injury). We observe clear positive correlations as assessed by a linear regression for $K$, $S_{norm}$, $C$, and $E$, and a clear negative correlation for $B$ (Table S 6). **D.** Mean community size for functional and spatial communities in sub-sampled multilayer networks networks (paired $t-test$, $t$ = 0.8896, $df$ = 22, $p$ = 0.3833). E Average module participation, the fraction of modules that contain at least one of that cell type, as determined based on spatial distance and functional connectivity for both randomly sub-sampled neurons and astrocytes (differences between groups are not statistically significant; ordinary one-way ANOVA, $F(3, 88)$ = 0.7745, $p$ = 0.5113.

| | $B$ | $C$ | $E$ |
|---|---|---|---|
| 95% CI of slope | [-0.1256,-0.02158] | [0.5094, 0.8444] | [0.5523, 0.8805] |
| $R^2$ | 0.3034 | 0.7803 | 0.8057 |
| $F$ | 8.709 | 71.05 | 82.94 |
| $DF$ | 20 | 20 | 20 |
| $p$ | 0.0079 | $<$0.0001 | $<$0.0001 |

**Supplementary Table 6.** Results of simple linear regression of $B$, $C$, and $E$ on network density $\kappa$ for sub-sampled multilayer networks at the final experimental time point.

| | $C$ | | | $B$ | | | $E$ | | |
|---|---|---|---|---|---|---|---|---|---|
| | $\beta$ | z | p | $\beta$ | z | p | $\beta$ | z | p |
| Intercept | 0.1589* | 4.702 | 0.000 | 0.0376* | 1.979 | 0.048 | 0.1192* | 5.709 | 0.000 |
| Strength | 0.7010* | 10.003 | 0.000 | -0.0641 | -1.627 | 0.104 | 0.8120* | 18.758 | 0.000 |
| MPEP | 0.0528 | 1.800 | 0.072 | 0.0177 | 1.074 | 0.283 | 0.0068 | 0.378 | 0.705 |
| Sham | -0.0047 | -0.151 | 0.888 | 0.0204 | 1.166 | 0.244 | -0.0194 | -1.011 | 0.312 |
| MPEP + Sham | 0.0229 | 0.509 | 0.611 | -0.0270 | -1.069 | 0.285 | 0.0164 | 0.589 | 0.556 |

**Supplementary Table 7.** Results of generalized linear regression to predict the effect of group assignment on mean clustering coefficient ($C$), mean normalized betweenness centrality ($B$), and global efficiency ($E$) for **sub-sampled multilayer** networks at the final experimental timepoint. $\beta$: estimated coefficient, $z$: value of test statistic for coefficient, the value of the estimate divided by the standard error of the estimate, and $p$: p-value for coefficient resulting from a t-test, pr($> z$), *df* = 17. The $z$-test tests the null hypothesis that the coefficient for that covariate is equal to zero. In this case, no topological measures were significantly affected by MPEP, injury, or their interaction.

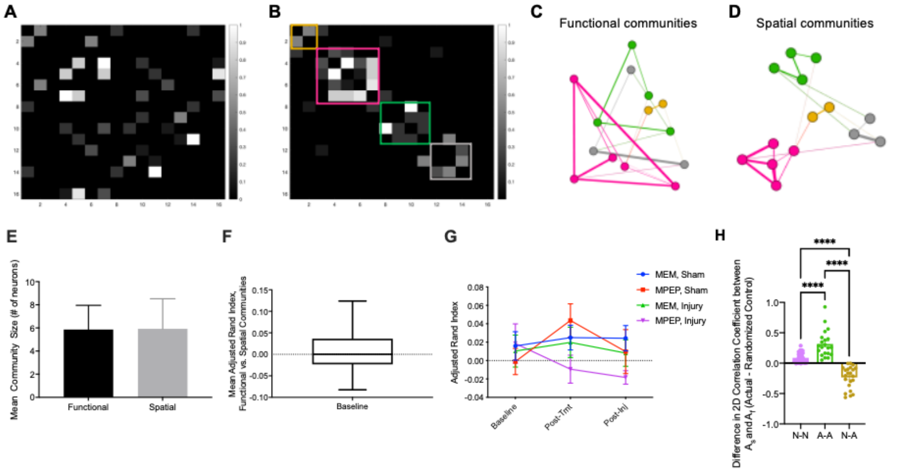


**Supplementary Figure 7.** Community structure of *in vitro* neuronal networks. **A.** Representative adjacency matrix of a neuronal network. **B.** The same adjacency matrix shown in panel **a** after community detection, reordered with modules grouped along the diagonal. **C.** Graph of the network shown in panel **A** and panel **B** depicting modularity as determined by functional connectivity between nodes. Nodes of the same color belong to the same functional community. **C.** Graph of the network shown in panel **A** and panel **B** depicting modularity as determined by spatial proximity between nodes. Nodes of the same color belong to the same spatial community. If functional connectivity were based on spatial proximity, the modules in panels **C** and **D** would be the same or similar. **E.** Number of functional (black) and spatial (gray) communities detected neuron networks (paired $t$-test, $t$=0.01640, $df$=35, $p$ = 0.9870). **F.** Mean Adjusted Rand Index for functionally- versus spatially-generated neuron-neuron networks. Whiskers range from the minimum value to the maximum value (95% CI of mean [-0.004606, 0.03082]). **G.** Adjusted Rand Index for functionally- versus spatially-generated neuron-neuron networks at the three measured time points for all four treatment groups (two-way ANOVA, time factor, $F(2, 96)$ = 0.7336, $p$=0.4828; treatment factor, $F(3, 96)$ = 1.404, $p$ =0.2463; interaction term, $F(6, 96)$ = 0.9866, $p$=0.4388). **H.** Difference in 2-D correlation coefficient between actual and randomized functional adjacency matrices and spatial adjacency matrices for neuron-neuron (purple), astrocyte-astrocyte (green), and neuron-astrocyte (brown) multilayer network layers. Error bars indicate standard error of the mean (SEM) and asterisks indicate statistical significance (no asterisks, ns, $*p \leq 0.05$, $**p \leq 0.01$, $***p \leq 0.001$, $****p \leq 0.0001$). Tmt: treatment; MEM: treated with minimum essential media; MPEP: treated with anti-mGluR$_5$; Injury: subjected to targeted neuronal tap injury; Sham: negative injury control.

| A. Neuron-Neuron | | | | |
|---|---|---|---|---|
| | MEM, Sham | MPEP, Sham | MEM, Inj | MPEP, Inj |
| 95% CI of slope | [0.011, 0.082] | [0.112, 0.174] | [0.204, 0.286] | [0.128, 0.163] |
| $R^2$ | 0 | 0.006 | 0.01 | 0.001 |
| F | 6.687 | 80.88 | 140.4 | 18.17 |
| DF | 15,086 | 14,159 | 14,001 | 22,879 |
| p | 0.00972 | 2.69E-19 | 3.12E-32 | 2.03E-05 |

| B. Astrocyte-Astrocyte | | | | |
|---|---|---|---|---|
| | MEM, Sham | MPEP, Sham | MEM, Inj | MPEP, Inj |
| 95% CI of slope | [0.691, 1.485] | [-.250, -0.005] | [0.652, 0.942] | [-0.066, 0.141] |
| $R^2$ | 0.075 | 0.002 | 0.096 | 0 |
| F | 29.07 | 4.185 | 129.4 | 0.4987 |
| DF | 359 | 1,674 | 1,221 | 2,197 |
| p | 1.27E-07 | 0.0409 | 1.45E-28 | 0.48 |

| C. Neuron-Astrocyte | | | | |
|---|---|---|---|---|
| | MEM, Sham | MPEP, Sham | MEM, Inj | MPEP, Inj |
| 95% CI of slope | [-0.191, -0.002] | [0.003, 0.091] | [0.142, 0.305] | [-0.042, 0.034] |
| $R^2$ | 0.001 | 0.001 | 0.007 | 0 |
| F | 4.019 | 4.358 | 28.97 | 0.03981 |
| DF | 3,792 | 7,009 | 4,417 | 11,344 |
| p | 0.0451 | 0.0369 | 7.75E-08 | 0.842 |

**Supplementary Table 8.** Results of linear regressions to predict functional adjacency from spatial adjacency for each experimental group at the final experimental time point (1 hour post-injury, see Table S2). Reported are the 95% confidence interval on the slope, the $R^2$ value, $F$-statistic, degrees of freedom ($DF$), and the $p$-value on the $F$-statistic for neuron-neuron networks (**A**), astrocyte-astrocyte networks (**B**) and multilayer networks (**C**). See Fig. X.

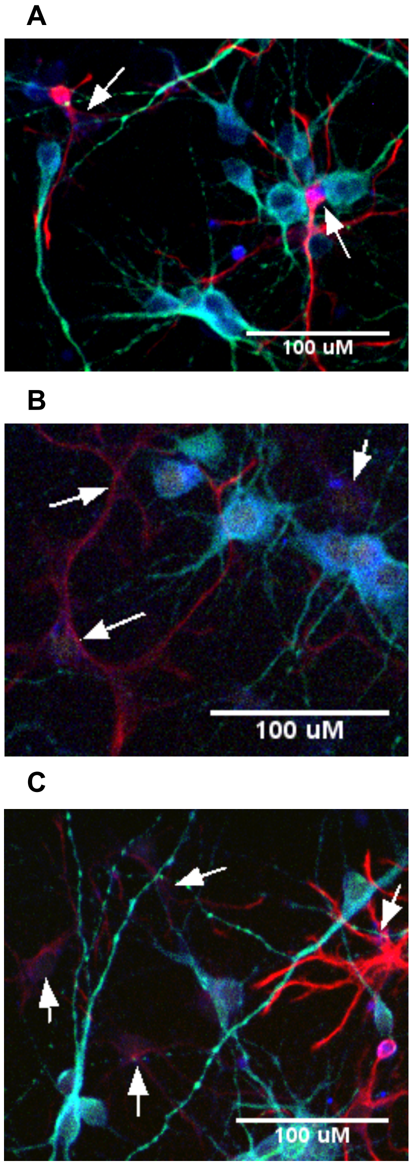


**Supplementary Figure 8.** Immunofluorescent staining confirms expression of $mGluR_5$ in neurons and astrocytes. Cells were stained for GFAP (red), an astrocytic marker, MAP2 (green), a neuronal marker, and $mGluR_5$ (blue). Turquoise and magenta areas represent co-localization of $mGluR_5$ on neurons and astrocytes, respectively. Shown in panels **a** - **c** are cropped fields of view from three dishes. Astrocytes expressing $mGluR_5$ are indicated with white arrows.

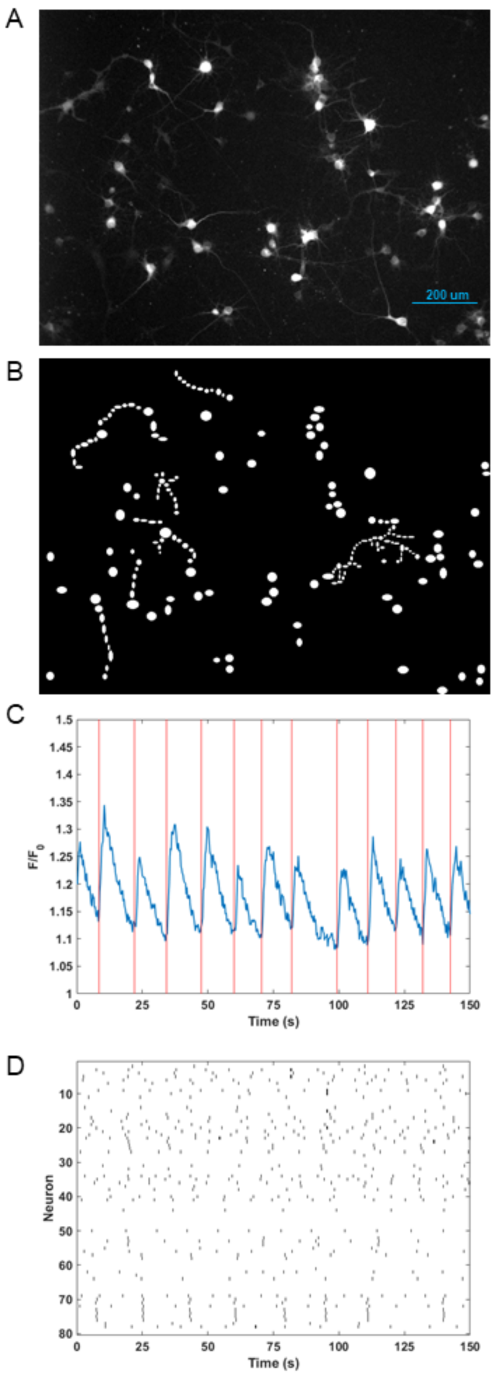


**Supplementary Figure 9.** *In vitro* calcium image acquisition and processing of neuron-astrocyte networks. **A.** Maximum fluorescence projection of the video recording. **B.** Manually identified ROIs segmenting neurons and astrocytes segments. **C.** Scaled fluorescence trace for a single neuronal ROI (blue), with detected spikes overlaid (vertical red lines). **D.** Raster plot of spikes over time, with each black vertical line indicating one spike, of all neuronal ROIs in the field of view.

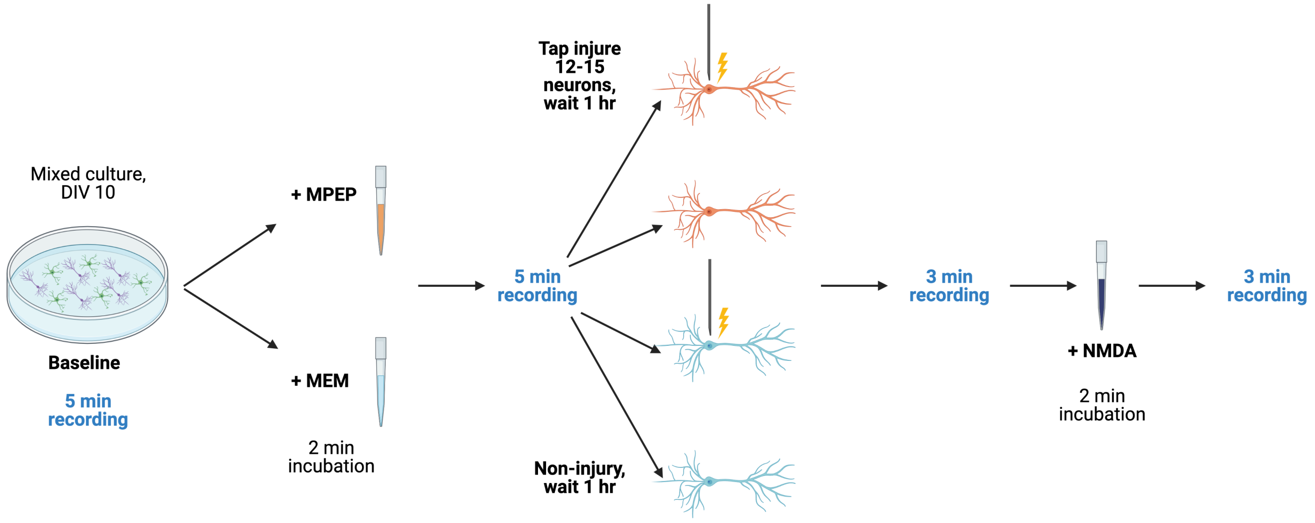


**Supplementary Figure 10.** Experimental design and treatment protocol (n = 8-9 dishes in each arm). All experiments were performed in mixed neuron (purple cells) and astrocyte (green cells) cultures at 10 days *in vitro* (DIV 10). Following a two-three minute equilibration period, five minutes of baseline calcium activity was recorded. 1uM MPEP HCl (orange) or MEM (light blue) was added, and five minutes of calcium activity was imaged following a two-minute incubation period. In half of the dishes, 12-15 neurons were mechanically injured via tap (indicated by yellow lightening bolt) with a pulled glass micropipette tip (black) controlled by a micro-manipulator (not shown). Three minutes of activity was imaged one hour later, followed by three minutes of imaging after addition of 100uM NMDA to differentiate neurons and astrocytes (see Methods). Renderings were created using Biorendering and are not to scale.

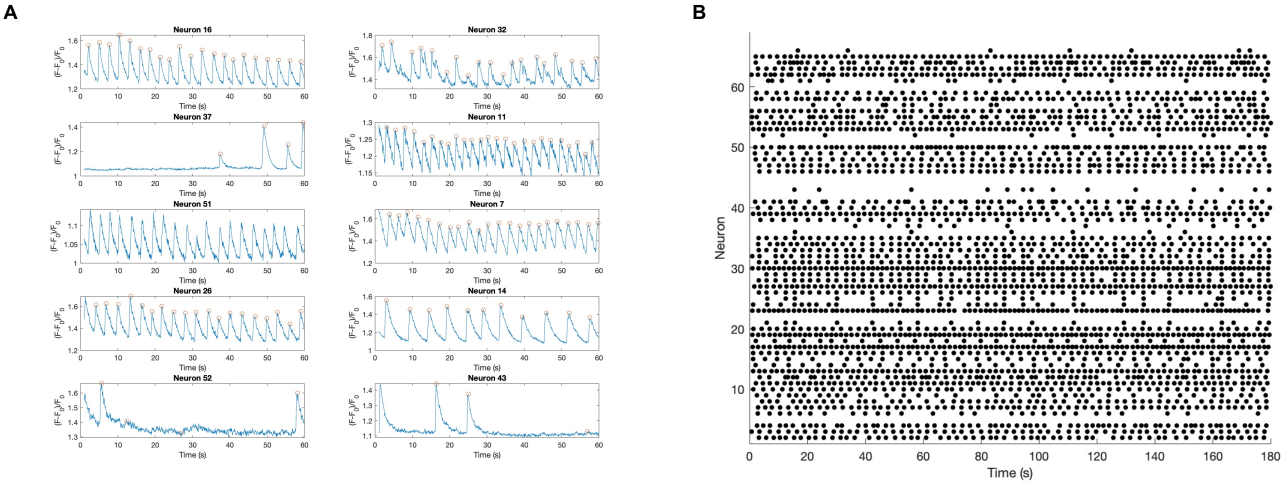


428 **Supplementary Figure 11.** Spike inference from neuronal calcium activity. **A.** Scaled calcium fluorescence traces of 10 randomly selected neurons from a
429 representative dish, with detected spikes (red open circles) overlaid. Activity is shown for the first minute of baseline recording. **B.** Raster plot showing spikes
430 over time for each neuron in the dish in **A** at baseline (entire recording). We note the absence of bursting activity characteristic of autaptic cultures.

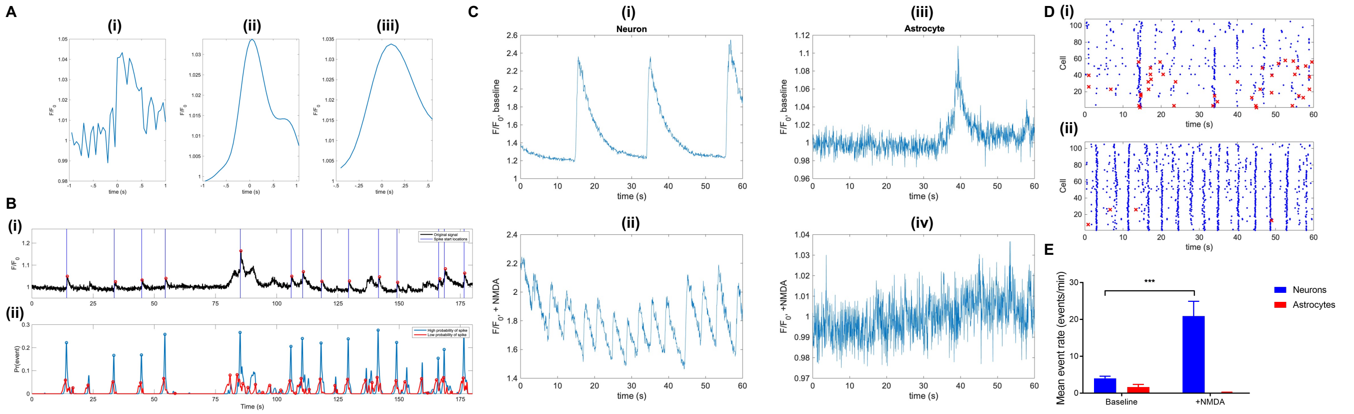


**Supplementary Figure 12.** Event inference from astrocyte calcium activity **A.** Processing of unfiltered astrocyte segment fluorescence data to generate a library waveform. **(i)** Unfiltered snippet of calcium activity around a peak. **(ii)** Filtered snippet of the same peak. **(iii)** Shortened snippet cropped to contain a single peak with shortened decay time, 75% of original duration. **B.** Predicted spikes generated by the automated astrocyte calcium event detection algorithm for an example astrocyte trace. **(i)** Scaled fluorescence of the astrocyte ROI with detected baseline (blue line) and peak (red circle) locations overlaid. **(ii)** High (red) and low (blue) probabilities of an event for the same trace, with detected peaks overlaid. The probability signals are used to determine event location. SNR = 1.7611. **C.** Scaled fluorescence and frequency of transients is higher after addition of NMDA in neurons but not astrocytes. Scaled fluorescence of a neuron before **(i)** and after **(ii)** addition of 100uM NMDA + 10uM glycine coagonist. Scaled fluorescence of an astrocyte before **(iii)** and after **(iv)** addition of NMDA. The astrocyte has fewer events after addition of NMDA. **D.** Raster plot showing population-level neuronal spiking (blue dots) and astrocytic calcium event (red crosses) activity before (**(i)**) and after (**(ii)**) addition of NMDA. Neuronal, but not astrocytic, population activity is visibly increased after the addition of NMDA. **E.** Mean neuronal (blue) and astrocytic (red) event rates at baseline (left) and following addition of 100uM NMDA + 1uM glycine coagonist (right). Addition of NMDA led to a significant increase in the number of events per minute for neurons (Sidak's multiple comparisons test; $p = 0.0003$, $t = 4.932$, $df = 8$), but not astrocytes (Sidak's multiple comparisons test; $p = 0.8165$, $t = 0.5897$, $df = 8$).

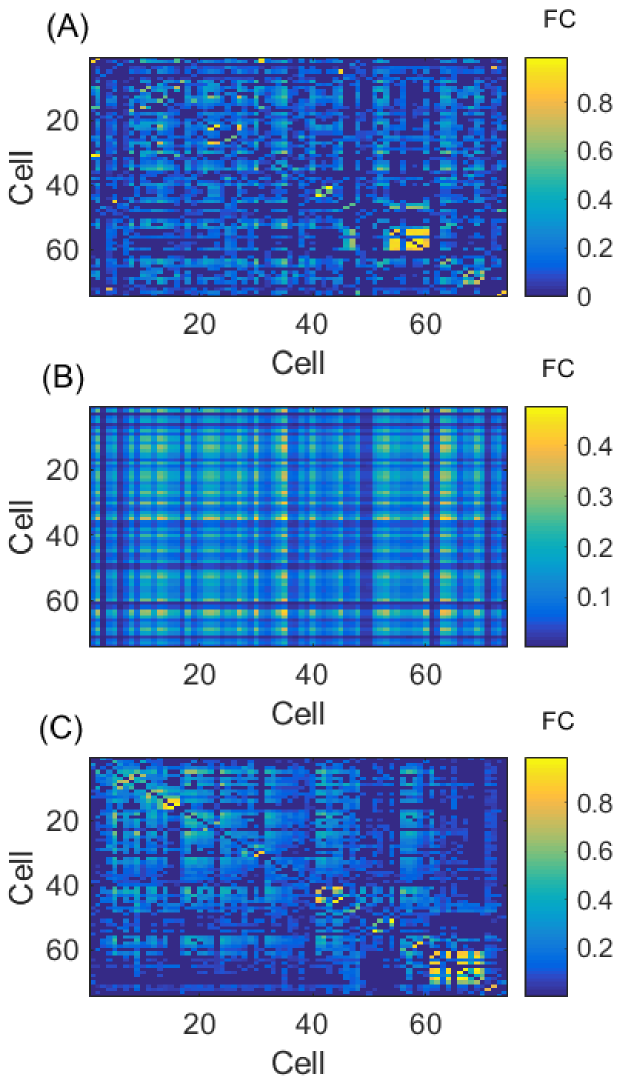


443 **Supplementary Figure 13.** Community detection in a multilayer functional network. **A.** Adjacency matrix of neurons and astrocyte segments based on
444 functional connectivity. **B.** Girvan-Newman null adjacency matrix for panel **A**. **C.** Adjacency matrix from panel **A** reordered by modular structure, so that
445 nodes in the same module are adjacent.
